# The pan-cancer lncRNA PLANE regulates an alternative splicing program to promote cancer pathogenesis

**DOI:** 10.1101/2020.12.04.412536

**Authors:** Liu Teng, Yu Chen Feng, Su Tang Guo, Pei Lin Wang, Shi Xing Wang, Sheng Nan Zhang, Teng Fei Qi, Ting La, Yuan Yuan Zhang, Xiao Hong Zhao, Didi Zhang, Jenny Y Wang, Yujie Shi, Jin Ming Li, Huixia Cao, Tao Liu, Rick F. Thorne, Lei Jin, Feng-Min Shao, Xu Dong Zhang

## Abstract

Genomic amplification of the distal portion of chromosome 3q, which encodes a number of oncogenic proteins, is one of the most frequent chromosomal abnormalities in malignancy. Here we functionally characterise a non-protein product of the 3q region, the long noncoding RNA (lncRNA) PLANE, which is upregulated in diverse cancer types through copy number gain as well as E2F1-mediated transcriptional activation. PLANE forms an RNA-RNA duplex with the nuclear receptor co-repressor 2 (NCOR2) pre-mRNA at intron 45, binds to heterogeneous ribonucleoprotein M (hnRNPM) and facilitates the association of hnRNPM with the intron, thus leading to repression of the alternative splicing (AS) event generating NCOR2-202, a major protein-coding NCOR2 AS variant. In consequence, PLANE promotes cancer cell proliferation and tumorigenicity and its upregulation is associated with poor patient outcomes. These results uncover the function and regulation of PLANE and suggest that PLANE may constitute a therapeutic target in the pan-cancer context.

## INTRODUCTION

Alternative splicing (AS) of precursor mRNAs (pre-mRNAs) is a fundamental mechanism that allows for the generation of diverse mature transcripts from a single gene thus amplifying the gene-coding capacity and increasing the functional diversity^1, 2, 3^. Over 95% of human multiexon genes undergo AS that is tightly controlled by the interaction of trans-acting proteins referred to as splicing factors with cis-acting nucleotide sequences^1, 2, 3^. Splicing factors encompass members of the serine-arginine (SR) protein family and heterogeneous ribonucleoproteins (hnRNPs) that promote or repress specific splicing events through interacting with exonic or intronic regulatory sequences classified as enhancers or silencers^1, 4^. Aberrant AS events are involved in the pathogenesis of many diseases including cancer through deregulating essential cellular processes such as cell survival and proliferation^1, 4, 5, 6^.

Nuclear receptor co-repressor 2 (NCOR2), also known as silencing mediator of retinoid and thyroid hormone receptors (SMRT) or T_3_ receptor-associating cofactor 1 (TRAC-1), acts as a central organising platform for assembling functional complexes which repress the transactivation of target genes. The NCOR2 N-terminal repression domains recruit other transcriptional corepressors such as histone deacetylases (HDACs) while its C-terminal interaction domains interact with nuclear receptors such as the thyroid hormone receptor and retinoic acid receptor^7, 8, 9^. Moreover, NCOR2-mediated repression also targets genes activated by other transcription factors such as AP-1 and NF-κB^9, 10, 11^. With its repression domains retained, NCOR2 exhibits varying affinities for different transcription factors through AS at its C-terminus^9, 12, 13^. Of note, cancer cells often display altered expression of NCOR2, implicating a role of its deregulation in cancer pathogenesis^14, 15, 16, 17, 18^. For example, NCOR2 is downregulated in multiple myeloma and and its low expression is associated the development of non-Hodgkin’s lymphoma and poor prognosis of lung adenocarcinoma (LUAD) patients^14, 15, 16, 17^. In contrast, high NCOR2 expression is linked to earlier recurrence of breast carcinoma (BRCA)^18^.

There is increasing appreciation of the role of long noncoding RNAs (lncRNAs) in cancer development and progression^1, 19, 20, 21, 22^. In particular, a growing number of lncRNAs have been linked to the deregulation of AS in cancer cells^23, 24, 25, 26, 27, 28^. LncRNAs regulate AS primarily through binding to splicing factors, associating with pre-mRNAs and impinging on chromatin remodelling^24^. For instance, the lncRNA MALAT1 regulates alternative splicing of a set of pre-mRNAs through modulating SR splicing factor phosphorylation and sub-nuclear localization, and is thus involved in the development, progression and treatment resistance of many types of cancers^26^, whereas the lncRNA LNIC01133 interacts with the SR splicing factor SRSF6 (SRp55) resulting in inhibition of epithelial-mesenchymal transition (EMT) and metastasis^27^. Moreover, the lncRNA SAF binds to the *Fas* pre-mRNA and recruits splicing factor 45 (SPF45) leading to generation of a Fas AS variant that protects cancer cells from Fas-induced cell death^28^.

Here we present evidence that the lncRNA PLANE forms an RNA-RNA duplex with the NCOR2 pre-mRNA and recruits hnRNPM, thus facilitating hnRNPM-mediated repression of the AS event generating NCOR2-202, a major protein-coding NCOR2 transcript variant. The resulting downregulation of NCOR2 at the protein level contributes to the increased proliferation and tumorigenicity of cancer cells. Moreover, we show that PLANE is frequently upregulated in diverse cancer types through genomic amplification and E2F1-mediated transcriptional activation, with practical implications of interference with PLANE as potential treatment approach in the pan-cancer context.

## RESULTS

### Genomic amplification and transcriptional activation by E2F1 drive PLANE upregulation in diverse cancer types

Through interrogating the lncRNA expression data in the Cancer Genome Atlas (TCGA)^29^, we identified a panel of eighteen pan-cancer upregulated lncRNAs that were increased in expression in at least 19 of 20 cancer types in relation to corresponding normal tissues (Supplementary Fig. 1a). Among them was melanotransferrin (MELTF, also known as MFI2) antisense RNA1 (MELTF-AS1 or MFI2-AS1) that is encoded by a gene located to the distal portion of chromosome 3q (3q29) (Supplementary Fig. 1a-c), whose amplification is one of the most prevalent chromosomal abnormalities observed in various cancer types^30, 31, 32, 33, 34^. Indeed, *MELTF-AS1* was the most frequently amplified gene among those that encode the pan-cancer upregulated lncRNAs (Supplementary Fig. 1d). We therefore sought to investigate the potential role of MELTF-AS1 in cancer pathogenesis. Of the five annotated MELTF-AS1 isoforms (Vega Genome Browser), the longest isoform was markedly more abundant than others in multiple cancer cell lines, including A549 and H1299 LUAD, NCI-H226 lung squamous cell carcinoma (LUSC), HCT116 colon adenocarcinoma (COAD), MCF-7 BRCA and Eca109 esophageal squamous carcinoma (ESCC) (Supplementary Fig. 1e, f). We hereafter focused on this isoform and renamed it PLANE (<u>P</u>an-cancer <u>L</u>ncRNA <u>A</u>ctivating <u>N</u>COR2 responsive to <u>E</u>2F1) given its functional relationship with NCOR2 and transcriptional responsiveness to E2F1 (see below). PLANE consists of 4 exons (E1-E4), with minimum free energy modelling predicting a broadly symmetrical structure with E1 and E3 constituting each pole, whereas E2 and E4 contributing to both poles of the molecule (Supplementary Fig. 1g).

We confirmed the cancer-associated upregulation of PLANE in cohorts of formalin-fixed paraffin-embedded (FFPE) LUSC, LUAD and COAD samples compared to paired adjacent normal tissues (Fig. 1a). Noticeably, despite the common increase in cancer tissues, PLANE levels did not differ among tumours of different stages (Supplementary Fig. 2a & Supplementary Tables 1-3). Likewise, there were no significant differences in PLANE expression between LUSC, LUAD and COAD of different groups stratified by tumour grade and patient gender as well as their median age at diagnosis (Supplementary Tables 1-3). Moreover, no significant changes were found in PLANE expression levels between COAD and colon adenomas (pre-neoplastic colon lesions), whereas PLANE expression was increased in colon adenomas compared with normal colon epithelia (Supplementary Fig. 2b). Collectively, these results suggest that PLANE upregulation is an early event during tumorigenesis. Furthermore, high PLANE expression was associated with poorer overall patient survival (OS) in diverse cancer types (Fig. 1b & Supplementary Fig. 2c), implicating its broad involvement in cancer development and progression.

**Figure 1.**
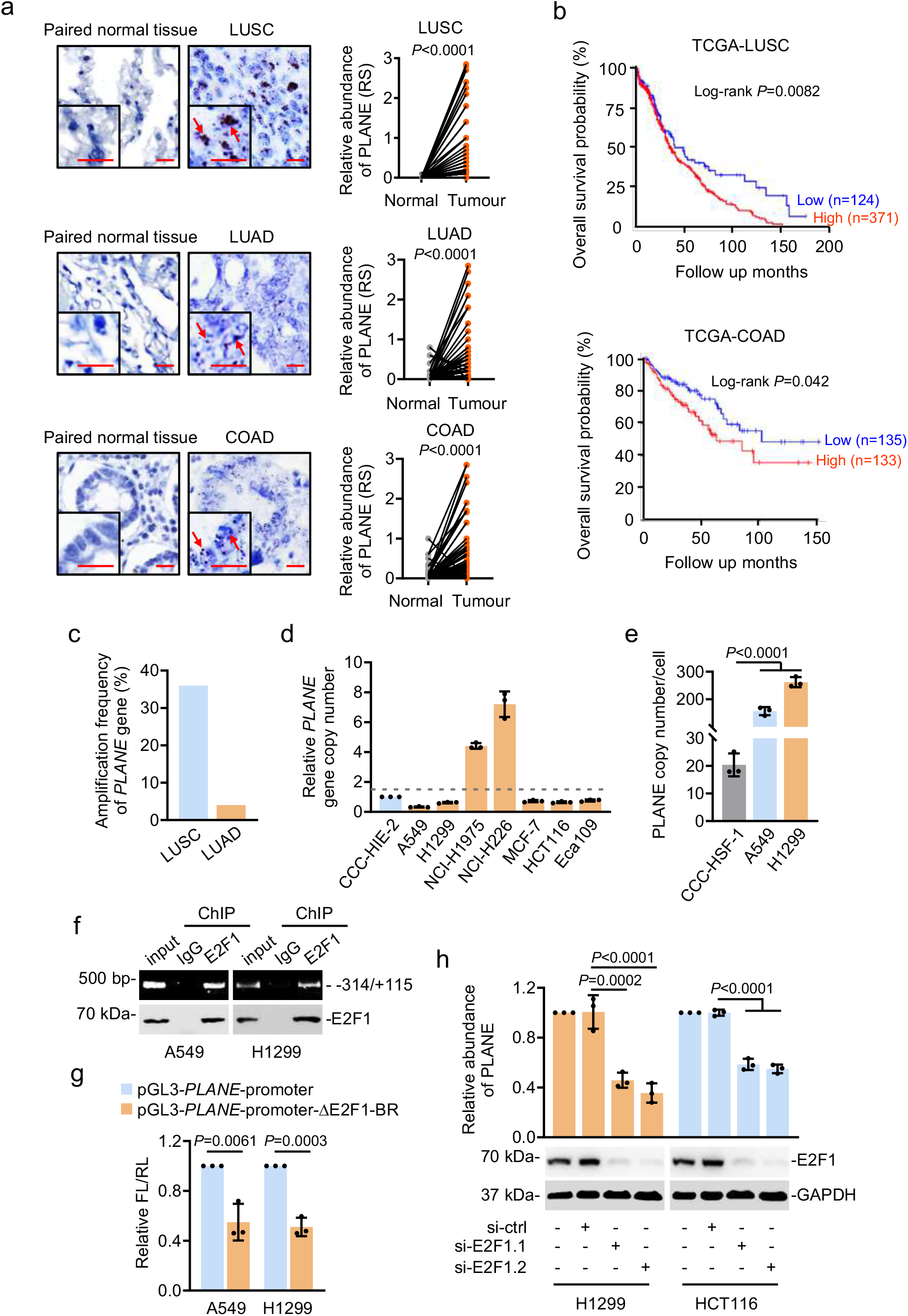
Genomic amplification and transcriptional activation by E2F1 drive PLANE expression that is upregulated in diverse cancer types. **a** Representative microscopic photographs of *in situ* hybridization (ISH) analysis of PLANE expression in formalin-fixed paraffin-embedded (FFPE) LUSC, LUAD and COAD tissues (n=75, 87 and 79 biologically independent samples, respectively) compared with corresponding paired adjacent normal tissues. Quantitation of PLANE expression in cancer relative to paired normal tissues is also shown. Scale bar, 5 μm. RS: reactive score. Two-tailed Student’s *t*-test. **b** Kaplan-Meier analysis of the probability of overall survival of LUSC (n = 495) and COAD (n = 268) patients derived from the TCGA datasets using the quartile (LUSC) or median (COAD) of PLANE levels as the cutoff. **c** qPCR analysis of genomic DNA from LUSC (n=22) and LUAD (n=24) tissues and corresponding paired adjacent normal tissues. A ≥1.5-fold increase in the copy number in the cancer tissue relative to the corresponding normal tissue is considered genomic amplification. **d** qPCR analysis of genomic DNA from the indicated cancer cell lines and the normal human intestinal epithelial cell line CCC-HIE-2. The copy number of *PLANE* in the CCC-HIE-2 cell line was arbitrarily designated as 1. A ≥1.5-fold increase in the copy number in cancer cell lines compared with the CCC-HIE-2 line is considered genomic amplification. Data are mean ± s.d.; n = 3 independent experiments. **e** Absolute quantitation of PLANE in A549 and H1299 cancer cells and normal human CCC-HSF-1 fibroblasts using qPCR. Data are mean ± s.d.; n = 3 independent experiments, one-way ANOVA followed by Tukey’s multiple comparisons test. **f** Chromatin immunoprecipitation (ChIP) analysis of the association between endogenous E2F1 and the region enriched of E2F1 binding motifs at the promoter of *PLANE* in A549 and H1299 cells. Data are representatives of 3 independent experiments. **g** The transcriptional activity of a *PLANE* reporter construct was reduced by deletion of the E2F1-binding region (E2F1-BR) at the promoter of *PLANE* in A549 and H1299 cells. Data are mean ± s.d.; n = 3 independent experiments, two-tailed Student’s *t*-test. FL: Firefly luciferase activity; RL: Renilla luciferase activity. **h** E2F1 silencing downregulated PLANE expression in H1299 and HCT116 cells. Data are representatives or mean ± s.d.; n = 3 independent experiments, one-way ANOVA followed by Tukey’s multiple comparisons test.

Consistent with the contribution of genomic amplification to the upregulation of PLANE in some cancer tissues (Supplementary Fig. 1d), qPCR analysis of genomic DNA demonstrated *PLANE* copy number gains in ~36% of LUSC (8 of 22) and ~4% of LUAD (1 of 24) (Fig. 1c). Increases in *PLANE* copy numbers were also evident in 2 of 7 cancer cell lines compared with the CCC-HIE-2 normal human intestinal epithelial cell line (Fig. 1d). Nonetheless, similar to the pan cancer upregulation of PLANE in tissue samples, all the cancer cell lines examined expressed higher levels of PLANE than the CCC-HSF-1 normal skin fibroblast cell line irrespective of their amplification status (Supplementary Fig. 2d), indicating that additional causal mechanisms such as transcriptional regulation is involved in the upregulation of PLANE in cancer cells. In support, absolute quantitation showed that there were respectively ~157 and ~262 PLANE molecules per A549 and H1299 cell, which did not have copy number gains in the *PLANE* gene, compared with ~20 PLANE molecules per CCC-HSF-1 cell (Fig. 1e).

To gain insights into transcriptional mechanisms involved in PLANE upregulation in cancer cells, we analysed its promoter for transcription factor binding sites using bioinformatics. This predicted multiple E2F1-binding motifs located to the −314/−14 region of the proximal promoter of the *PLANE* gene (Supplementary Fig. 3a)^35^. Indeed, this region (E2F1-BR) was co-precipitated with endogenous E2F1 and was required for transcriptional upregulation of PLANE as the transcriptional activity of a *PLANE* reporter construct was inhibited when the E2F1-BR was deleted (Fig. 1f, g). Moreover, co-transfection of E2F1 selectively enhanced the transcriptional activity of the *PLANE* reporter whereas knockdown of E2F1 diminished reporter activity (Supplementary Fig. 3b, c), supporting the notion that *PLANE* is transcriptionally activated by E2F1 through the identified E2F1-BR. In accordance, knockdown of E2F1 reduced the endogenous PLANE levels, whereas overexpression of E2F1 caused PLANE upregulation (Fig. 1h & Supplementary Fig. 3d). Furthermore, PLANE levels were correlated with E2F1 expression levels in diverse cancer types (Supplementary Fig. 3e). Collectively, these results demonstrate that E2F1 along with genomic amplification are responsible for the upregulation of PLANE in cancer cells.

The gene encoding PLANE is divergently located opposite to the protein-coding gene *MELTF* (Supplementary Fig. 4a). Nevertheless, knockdown of PLANE did not impinge upon MELTF expression (Supplementary Fig. 4b), and similarly, knockdown of MELTF did not affect PLANE expression levels (Supplementary Fig. 4c). Thus, there is no regulatory interaction between PLANE and its neighbouring gene *MELTF*.

### PLANE promotes cancer cell proliferation and tumorigenicity

We examined the biological significance of PLANE upregulation in cancer cells. SiRNA knockdown of PLANE inhibited cell proliferation and reduced clonogenicity in diverse cancer cell lines (Fig. 2a-c), which was associated with G0/G1 cell cycle arrest (Supplementary Fig. 5a). Conversely, overexpression of PLANE increased, albeit moderately, proliferation in A549 and H1299 cells (Supplementary Fig. 5b). Gene set enrichment analysis (GSEA) of the RNA-sequencing (RNA-seq) data from A549 cells revealed that knockdown of PLANE caused downregulation of numerous genes of signalling pathways involved in cell cycle progression, including the E2F1, G2/M checkpoint and mitotic spindle assembly pathways (Supplementary Fig. 5c). Thus, PLANE expression promotes the integral proliferative machinery of cancer cells.

**Figure 2.**
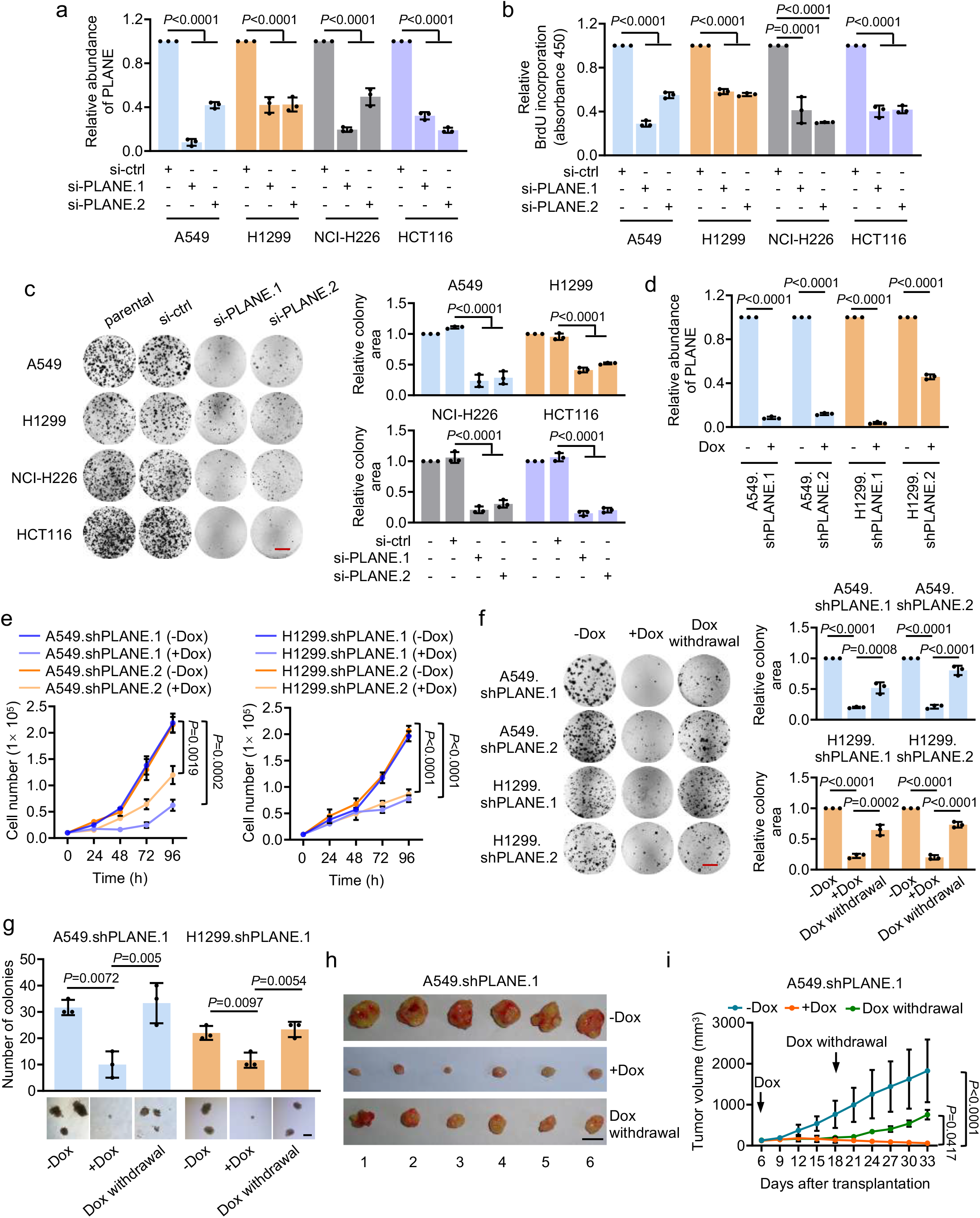
PLANE promotes cancer cell proliferation and tumorigenicity. **a-c** SiRNA knockdown of PLANE (a) inhibited 5-bromo-2’-deoxyuridine (BrdU) incorporation (b) and clonogenicity (c) in multiple cancer cell lines. Relative clonogenicity was quantitated using ImageJ-plugin ‘ColonyArea’. Data are mean ± s.d. or representatives; n = 3 independent experiments, two-tailed Student’s *t*-test. Scale bar, 1 cm. **d** Induced knockdown of PLANE by the addition of doxycycline (Dox, 500 nM) in A549.shPLANE and H1299.shPLANE cells. Data are mean ± s.d.; n = 3 independent experiments, two-tailed Student’s *t*-test. **e** Doxycycline (Dox, 500nM)-induced knockdown of PLANE inhibited A549.shPLANE and H1299.shPLANE cell proliferation as shown by decelerated cell number increases. Data are mean ± s.d.; n = 3 independent experiments, two-tailed Student’s *t*-test. **f** Induced knockdown of PLANE inhibited A549.shPLANE and H1299.shPLANE cell clonogenicity, which was partially reversed by cession of Dox treatment. Relative clonogenicity of cells was quantitated using ImageJ-plugin ‘ColonyArea’. Data are representatives or mean ± s.d.; n = 3 independent experiments, one-way ANOVA followed by Tukey’s multiple comparisons test. Scale bar, 1 cm. **g.** Representative microscopic photographs of anchorage-independent growth of A549.shPLANE and H1299.shPLANE cells with or without treatment with Dox and cessation of Dox treatment. Quantification of anchorage-independent growth of the cells is also shown. Scale bar, 0.5 mm. Data are representatives or mean ± s.d.; n = 3 independent experiments, one-way ANOVA followed by Tukey’s multiple comparisons test. **h & i** Representative photographs (h) and growth curves (i) of A549.shPLANE xenografts in nu/nu mice with or without treatment with Dox (2 mg/ml supplemented with 10 mg/ml sucrose in drinking water) and cessation of Dox treatment. Data are representatives or mean ± s.d.; n = 6 mice per group, one-way ANOVA followed by Tukey’s multiple comparison test. DOX: 2 mg/ml supplemented with 10 mg/ml sucrose in drinking water.

To facilitate further investigations, we established A549 and H1299 sublines (A549.shPLANE and H1299.shPLANE) with conditional knockdown of PLANE in response to doxycycline (Dox) (Fig. 2d). Induced PLANE knockdown similarly triggered reductions in cell proliferation and clonogenicity and induced anchorage-independent growth in A549.shPLANE and H1299.shPLANE sublines (Fig. 2e-g). Moreover, Dox treatment of nu/nu mice retarded the growth of A549.shPLANE.1 xenografts (Fig. 2h, i & Supplementary Fig. 5d, e). Cessation of Dox treatment restored the expression of PLANE and recovered, at least in part, the clonogenic potential *in vitro* and tumour xenograft growth in mice (Fig. 2h, i & Supplementary Fig. 5d, e), further consolidating the role of PLANE in tumorigenicity.

### PLANE regulates NCOR2 pre-mRNA AS

To dissect the mechanisms whereby PLANE promotes cancer cell proliferation, we compared the transcriptomes of A549 cells with and without PLANE knockdown. The results showed that the NCOR2 AS variant, NCOR2-202 (ENST00000397355.1; grch37.ensembl.org), was the most highly upregulated transcript associated with knockdown of PLANE (Fig. 3a). Interestingly, there were no significant changes in the levels of other protein-coding NCOR2 AS variants, including NCOR2-001 and NCOR2-005, which along with NCOR2-202, give rise to the three major NCOR2 protein isoforms, NCOR2-3, −1 and −2, respectively (Fig. 3a & Supplementary Fig. 6a)^36^. Due to sequence overlaps it was not feasible to specifically confirm the increase in the NCOR2-202 AS variant using qPCR (Supplementary Fig. 6a), but analysis using primers recognising a common region present in NCOR2-001, NCOR2-202 and NCOR2-005 demonstrated increased expression following PLANE knockdown (Fig. 3b & Supplementary Fig. 6a), conceivably reflecting NCOR2-202 upregulation. Furthermore, immunoblotting with NCOR2 antibodies against residues near its N-terminus that are conserved in NCOR2 protein isoforms 1-3 demonstrated that knockdown of PLANE caused an increase in NCOR2 protein expression (Fig. 3b & Supplementary Fig. 6a), suggesting that the increase in the NCOR2-202 AS variant was translated to the upregulation of the NCOR2 protein. Consistently, overexpression of PLANE caused downregulation of NCOR2 protein levels (Fig. 3c).

**Figure 3.**
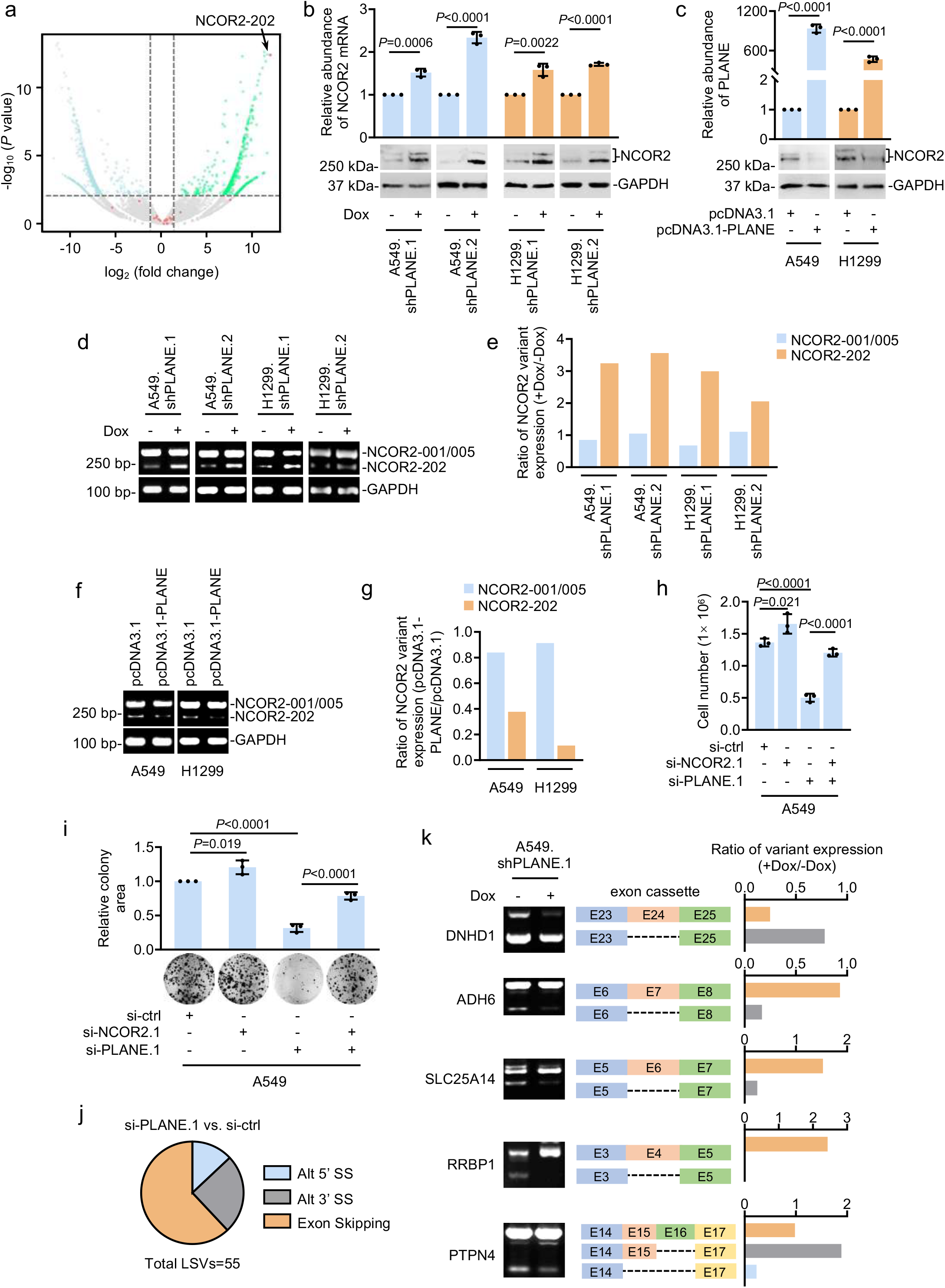
PLANE represses NCOR2-202-generating AS event. **a** Volcano plot of transcript expression derived from RNA-seq data showing that the NCOR2 AS variant NCOR2-202 was the most upregulated transcript and the only NCOR2 AS variant that was increased in A549 cells caused by siRNA knockdown of PLANE. Red dots represent NCOR2 AS variants. n = 2 experimental repeats. **b** qPCR analysis using primers spanning across a common region present in NCOR2-001, NCOR2-202 and NCOR2-005 and Western blotting analysis using an anti-NCOR2 antibody against residues near its N-terminus that is conserved in NCOR2 protein isoform NCOR2-3, −1 and −2 encoded individually by NCOR2-001, NCOR2-202 and NCOR2-005 showing upregulation of NCOR2 at the mRNA and protein levels, respectively, by induced knockdown of PLANE in A549.shPLANE and H1299.shPLANE cells. Data are representatives or mean ± s.d.; n = 3 independent experiments, two-tailed Student’s *t*-test. **c** Overexpression of PLANE caused downregulation of NCOR2 at the protein level in A549 and H1299 cells. Data are representatives or mean ± s.d.; n = 3 independent experiments, two-tailed Student’s *t*-test. **d** Inducible knockdown of PLANE promoted the NCOR2-202-generating AS event but did not affect the AS event giving rise to NCOR2-001 and NCOR2-005 as shown in RT-PCR analysis. Data are representatives of 3 independent experiments. **e** Relative levels of NCOR2-202/-001/-005 in cells with or without induced knockdown of PLANE as shown in **d**. Data are representatives of 3 independent experiments. **f** Overexpression of PLANE reduced the NCOR2-202-generating AS event but did not affect the AS event giving rise to NCOR2-005 as shown in RT-PCR analysis. **g** Relative levels of NCOR2-202/-001/-005 in cells with or without overexpression of PLANE as shown in **f**. Data are representatives of 3 independent experiments. **h & i** Co-knockdown of NCOR2 using siRNA partially reversed siRNA-knockdown of PLANE-induced inhibition of A549 cell proliferation as shown by decelerated cell number increases (h) and clonogenicity (i). Relative clonogenicity of cells was quantitated using ImageJ-plugin ‘ColonyArea’. Data are representatives or mean ± s.d.; n = 3 independent experiments, one-way ANOVA followed by Tukey’s multiple comparison test. **j** MAJIQ analysis of RNA-seq data (two experimental repeats) showing the categorization of alternative splicing events caused by PLANE knockdown in A549 cells. LSVs, local splicing variations; Alt 5’ SS, Alternative 5’ splicing site; Alt 3’ SS, Alternative 3’ splicing site. MAJIQ, modelling alternative junction inclusion quantification. **k** RT-PCR analysis of the indicated AS events using primers flanking PLANE-regulated alternative exons. Relative levels of relevant AS variants quantitated using densitometry are also shown. Data are representatives of 3 independent experiments. DNHD1, Dynein Heavy Chain Domain 1; ADH6, Alcohol Dehydrogenase 6; SLC25A14, Solute Carrier Family 25 Member 14; RRBP1, Ribosome Binding Protein 1; PTPN4, Protein Tyrosine Phosphatase Non-Receptor Type 4.

To verify that the upregulation of the NCOR2-202 AS variant caused by PLANE knockdown was due to a posttranscriptional increase, we tested the levels of the NCOR2 pre-mRNA in cells with and without knockdown of PLANE. The results showed that PLANE knockdown did not alter NCOR2 pre-mRNA expression (Supplementary Fig. 6b). Similarly, overexpression of PLANE did not cause any changes in NCOR2 pre-mRNA levels (Supplementary Fig. 6c). Moreover, knockdown of PLANE did not affect the enrichment of the transcriptional activation mark H3K4me3 and the transcriptional repression mark H3K27me3 to the *PLANE* promoter (Supplementary Fig. 6d)^37^. Collectively, these results point to posttranscriptional regulation of NCOR2 expression by PLANE through controlling the AS event generating the NCOR2-202 transcript. To substantiate this, we carried out semiquantitative RT-PCR analysis using primers flanking the splice site at the junction of exon/intron 45 which generates NCOR2-202. Detection of the splicing event generating NCOR2-001 and NCOR2-005 at the junction was included as a control. Here NCOR2-202 differs from NCOR2-001 and NCOR2-005 in its generation through an alternative 5’ splice site (Supplementary Figs. 6a & 7a). Instructively, PLANE knockdown resulted in an increase in the NCOR2-202-generating AS event but did not affect the event giving rise to NCOR2-001/NCOR2-005 (Fig. 3d, e). In contrast, overexpression of PLANE reduced the NCOR2-202-generating AS event (Fig. 3f, g). Thus, PLANE represses the AS event that produces NCOR2-202.

To investigate the biological impact of PLANE regulation of AS production of NCOR2-202, we tested the effect of siRNAs targeting common regions of NCOR2-001, NCOR2-005 and NCOR2-202 on inhibition of cell proliferation caused by PLANE knockdown (Supplementary Fig. 6a), which conceivably reflected the consequence of inhibition NCOR2-202 expression, as NCOR2 protein upregulation in PLANE knockdown cells was exclusively caused by the increase in NCOR2-202 (Fig. 3a, b & Supplementary Fig. 6a). For simplicity, we hereafter refer to these siRNAs as NCOR2 siRNAs. As anticipated, introduction of the NCOR2 siRNAs diminished the upregulation of NCOR2 and reversed, at least partially, the reduction in cell proliferation and clonogenicity caused by knockdown of PLANE (Fig. 3h, i & Supplementary Fig. 7b). Of note, NCOR2 knockdown alone caused moderate increases in cell proliferation (Fig. 3h, i & Supplementary Fig. 7b).

We also tested whether PLANE impinges on AS of other pre-mRNAs through analysing RNA-seq data from A549 cells using the modelling alternative junction inclusion quantification (MAJIQ) that detects and quantifies local splicing variations (LSVs), including classical binary splicing events and non-classical binary splits and splits involving more than two junctions^38^. MAJIQ analysis identified 46916 and 48635 AS events across 10187 genes in cells with and without PLANE knockdown, respectively, with 55 significant LSVs apart from the NCOR2-202-generating splicing event triggered by knockdown of PLANE (delta PSI ≥20%; confidence threshold >95%) (Fig. 3j). By use of semiquantitative RT-PCR we validated 5 randomly selected LSVs caused by knockdown of PLANE (Fig. 3k). Together, these results demonstrate that PLANE regulates AS of many other pre-mRNAs in addition to NCOR2, although its role in promoting cell proliferation is largely attributable to its modulatory effects on NCOR2 pre-mRNA AS.

### PLANE forms an RNA-RNA duplex with the NCOR2 pre-mRNA

PLANE predominantly localized to the nucleus as shown by ISH analysis of A549 cells grown on coverslips and qPCR analysis of subcellular fractions (Fig. 4a, b), suggesting the possibility through forming RNA-RNA duplexes with pre-mRNAs to regulate AS. Bioinformatics analysis using the IntaRNA program (http://rna.informatik.uni-freiburg.de) identified a potential PLANE-binding region (PLANE-BR) at intron 45 of the NCOR2 pre-mRNA that complements to a fragment enriched of duplex-forming oligonucleotides (DFOs) contained in PLANE (Supplementary Fig. 7c)^39^. To test whether PLANE forms RNA-RNA duplexes with the NCOR2 pre-mRNA, we employed a cell-free assay system. In vitro-synthesized biotin-labelled PLANE precipitated an RNA fragment containing the PLANE-BR at intron 45 of the NCOR2 pre-mRNA (Fig. 4c). However, this association was diminished when the PLANE-BR or the DFOs within PLANE were deleted (Fig. 4c). Moreover, biotin-labelled PLANE failed to precipitate a fragment of the NCOR2 pre-mRNA that did not contain the PLANE-BR (Fig. 4d). Consistently, in vitro-synthesized biotin-labelled PLANE also precipitated the NCOR2 pre-mRNA from A549 and H1299 cell nuclear extracts (Fig. 4e). In addition, endogenous PLANE bound to the PLANE-BR but not a non-PLANE-BR-containing fragment of endogenous NCOR2 pre-mRNA as shown in domain-specific chromatin isolation by RNA purification (dChIRP) assays (Fig. 4f). Collectively, these results reveal the formation of an RNA-RNA duplex between PLANE and the NCOR2 prem-mRNA through the DFOs and PLANE-BR, respectively. Of note, treatment of nuclear extracts from A549 cells with proteinase K did not disrupt the RNA-RNA duplex formed by PLANE and the NCOR2 pre-mRNA (Fig. 4g), demonstrating the binding between PLANE and the NCOR2 pre-mRNA is direct and not protein dependent.

**Figure 4.**
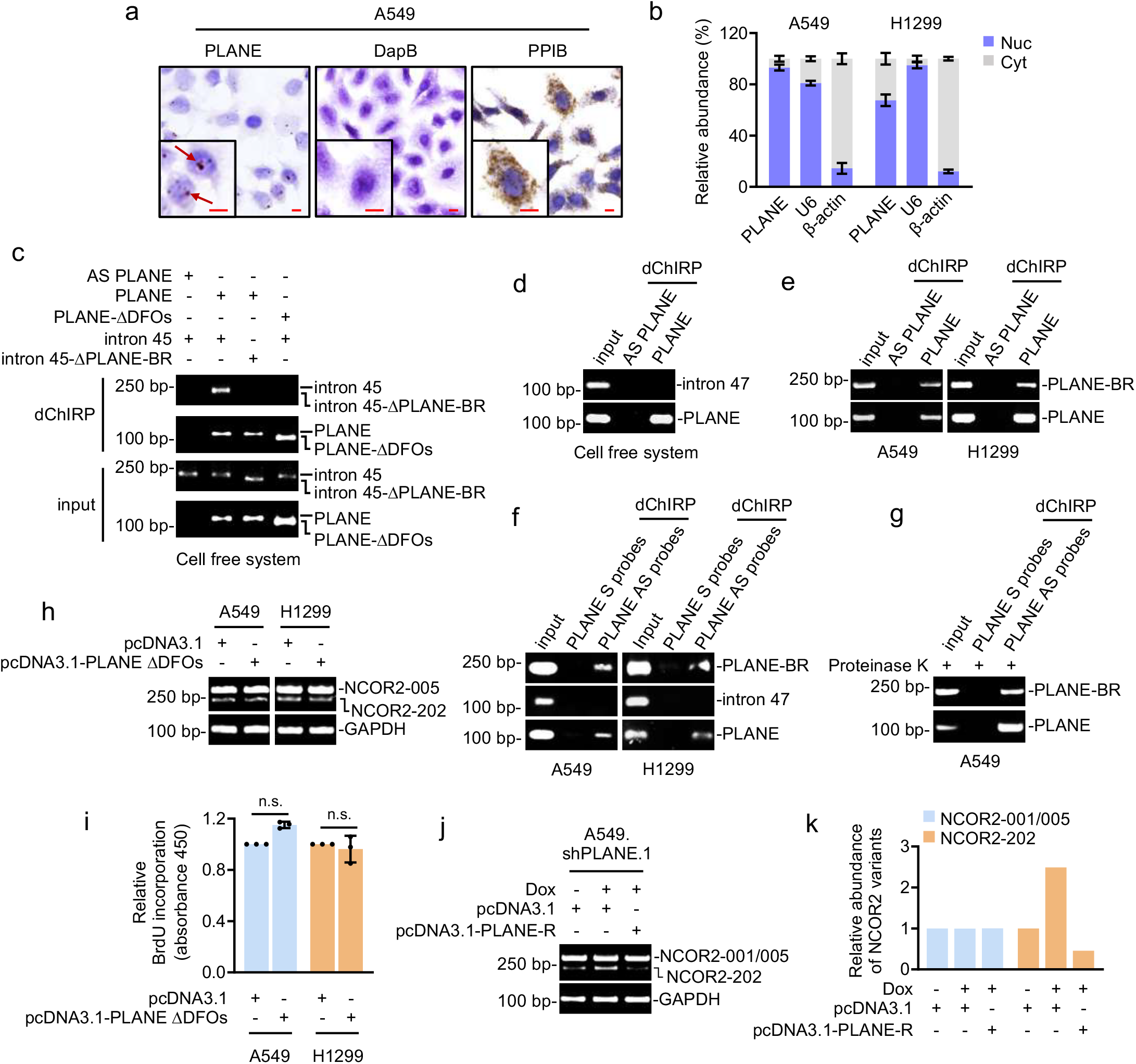
PLANE forms an RNA-RNA duplex with the NCOR2 pre-mRNA. **a** Representative microphotographs of *in situ* hybridization (ISH) analysis of PLANE expression in A549 cells grown on coverslips. Analysis of DapB and PPIB RNA expression was included as a negative and a positive control, respectively. Scale bar, 10 μm. Data are representatives of 3 independent experiments. **b** qPCR analysis of PLANE expression in the nuclear and cytoplasmic fractions of A549 and H1299 cells. Analysis of U6 and β-actin RNA expression was included as controls. Data are mean ± s.d.; n = 3 independent experiments, two-tailed Student’s *t*-test. Cyt: cytoplasm; Nuc: nucleus. **c** *In vitro*-synthesized biotin-labelled PLANE bound to the PLANE binding region (PLANE-BR) of *in vitro*-transcribed intron 45 of NCOR2 pre-mRNA as shown in domain-specific chromatin isolation by RNA purification (dChIRP) assays. This binding was abolished when the PLANE-BR at NCOR2 pre-mRNA or the duplex-forming oligonucleotides (DFOs) within PLANE were deleted (intron 45-ΔPLANE-BR and PLANE-ΔDFOs, respectively). Data are representatives of 3 independent experiments. **d** *In vitro*-transcribed biotin-labelled PLANE did not precipitate *in vitro*-transcribed intron 47 of the NCOR2 pre-mRNA that does not contain the PLANE-BR as shown in dChIRP assays. Data are representatives of 3 independent experiments. AS, antisense. **e** *In vitro*-synthesized biotin-labelled PLANE precipitated the endogenous NCOR2 pre-mRNA in A549 and H1299 cell nuclear extracts as shown in dChIRP assays. Data are representatives of 3 independent experiments. **f** Endogenous PLANE coprecipitated a fragment of the endogenous NCOR2 pre-mRNA containing the intact PLANE-BR but not a non-PLANE-BR-containing fragment (intron 47 of NCOR2 pre-mRNA) as shown in dChIRP assays. Data are representatives of 3 independent experiments. S, sense; AS, antisense. **g** PLANE co-precipitated a fragment of intron 45 of the NCOR2 pre-mRNA containing the PLANE-BR in A549 cells treated with proteinase K as shown in dChIRP assays. Data are representatives of 3 independent experiments. **h & i** Expression of a PLANE mutant with the DFOs deleted (PLANE-ΔDFOs) did not affect the NCOR2-202-generating splicing event (h) and 5-bromo-2’-deoxyuridine (BrdU) incorporation (i). Data are representatives or mean ± s.d.; n = 3 independent experiments, two-tailed Student’s *t*-test. **j** Expression of a shRNA-resistant PLANE mutant (PLANE-R) diminished the enhancement of the NCOR2-202-generating AS event caused by induced knockdown of PLANE in A549.shPLANE.1 cells. Data are representatives of 3 independent experiments. **k** Relative levels of NCOR2-202/-001-/005 as shown in **j** quantitated using densitometry. Data are representatives of 3 independent experiments.

We then examined the functional significance of the RNA-RNA duplex in PLANE-mediated regulation of NCOR2 pre-mRNA AS and cell proliferation. In contrast to overexpression of wild-type PLANE (Fig. 3f, g & Supplementary Fig. 5b), introduction of a PLANE mutant lacking the DFOs into A549 and H1299 cells had no effect on the NCOR2-202-generating splicing event and cell proliferation (Fig. 4h, i & Supplementary Fig. 7d). Moreover, introduction of a shRNA-resistant PLANE mutant (PLANE-R) inhibited the AS event caused by PLANE knockdown (Fig. 4j, k). Taken together with preceding data (Fig. 4c-k & Supplementary Fig. 7d), these findings indicate that the formation of the RNA-RNA duplex is required for the PLANE effects on NCOR2 pre-mRNA AS and cell proliferation.

### PLANE interacts with hnRNPM

We also interrogated the proteins that interact with PLANE using RNA-pulldown followed by mass spectrometry. The most abundant protein that coprecipitated with PLANE was hnRNPM (Fig. 5a & Supplementary Table 4), one of the hnRNP proteins that complex with heterogeneous nuclear RNA and are essential in regulating mRNA maturation processes including pre-mRNA splicing^40, 41^. The association between PLANE and hnRNPM was readily confirmed using RNA pulldown and RNA immunoprecipitation (RIP) assays (Fig. 5b, c). In contrast, no association was detected between PLANE and hnRNPK that was included as a control (Fig. 5b). Similarly, there was no association between hnRNPM and the mitochondrial lncRNA lncCyt b included as an additional control (Fig. 5c). In support of the direct interaction between PLANE and hnRNPM, *in vitro*-synthesized PLANE co-precipitated recombinant hnRNPM in a cell free system (Fig. 5d).

**Figure 5.**
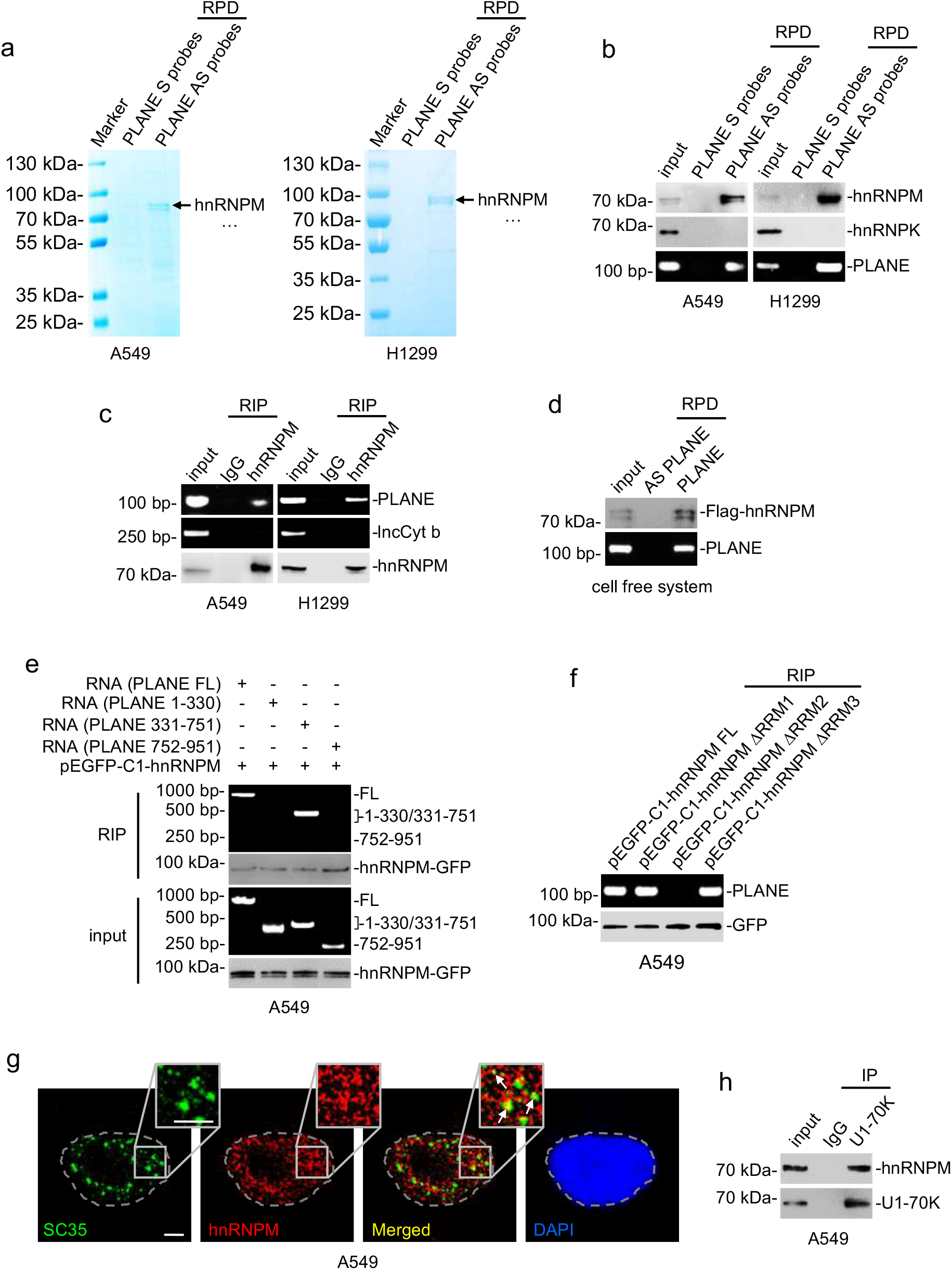
PLANE interacts with hnRNPM. **a** RNA pulldown followed by mass spectrometry analysis identified that hnRNPM is the most abundant protein co-pulled down with PLANE antisense probes in A549 and H1299 cells. S: sense; AS: antisense. n = 1 experiment. **b** hnRNPM was co-pulled down with PLNAE in A549 and H1299 cells as shown in RNA pulldown assays. hnRNPK was included as a negative control. S, sense; AS, antisense. Data are representatives of 3 independent experiments. **c** PLANE was coprecipitated with hnRNPM in A549 and H1299 cells as shown in RNA immunoprecipitation (RIP) assays. The lncRNA lncCyt b was included as a negative control. Data are representatives of 3 independent experiments. **d** Recombinant Flag-tagged hnRNPM was co-pulled down with *in vitro*-synthesized biotin-labelled PLANE as shown in RNA pulldown assays. Data are representatives of 3 independent experiments. **e** *In vitro*-synthesized full length (FL) PLANE and PLANE fragment 331-751 but not 1-330 or 752-951 were coprecipitated with hnRNPM as shown in RIP assays. Data are representatives of 3 independent experiments. **f** PLANE was co-precipitated with full-length (FL) hnRNPM, hnRNPM Δ RNA recognition motif (RRM) 1 and hnRNPM ΔRRM3 but not hnRNPM ΔRRM2 as shown in RIP assays. Data are representatives of 3 independent experiments. **g** Representative microscopic photographs of immunofluorescence staining showing co-localization of hnRNPM and SC35 in A549 cells grown on coverslips. Data shown are representatives of 3 independent experiments. Scale bar: 2 μm **h** hnRNPM was co-precipitated with U1-70K in A549 and H1299 cells. Data are representatives of 3 independent experiments. IP: immunoprecipitation.

To define the region of PLANE responsible for its interaction with hnRNPM, we carried out mapping experiments with PLANE mutants transcribed *in vitro* (Supplementary Fig. 8a). This analysis showed that PLANE fragment 331-751 but not fragment 1-330 or 752-951 was coprecipitated with hnRNPM (Fig. 5e), indicating that the 331-751 region of PLANE is required for its association with hnRNPM. We also conducted mapping experiments to identify the structural determinant for the binding of hnRNPM with PLANE. hnRNPM contains three RNA recognition motifs (RRMs) that are located at aa 72-147, aa 206-279, and aa 654-730, respectively (Supplementary Fig. 8b). Deletion of the aa 206-279 RRM but not the aa 72-147 or aa 206-279 RRM diminished the association between hnRNPM and PLANE (Fig. 5f), indicating that the aa 206-279 RRM of hnRNPM is necessary for its binding to PLANE.

### PLANE links hnRNPM to regulation of NCOR2 pre-mRNA AS

We next investigated the relationship of PLANE and hnRNPM in regulating NCOR2 pre-mRNA AS. As anticipated, hnRNPM predominantly localised to the nucleus in A549 and H1299 cells (Supplementary Fig. 8c). Noticeably, while a proportion of hnRNPM colocalized with the splicing factor SC35, a marker of nuclear speckles where the pre-mRNA splicing machinery is assembled, modified and stored (Fig. 5g)^42^, hnRNPM was also co-precipitated with U1 small nuclear ribonucleoprotein 70kDa (snRNP70; as known as U1-70K) that associates with the spliceosome small nuclear RNA (snRNA) U1 and is commonly used as a marker of spliceosomes (Fig. 5h)^25, 43^, consistent with its role as a splicing factor.

Through in silico analysis we identified a fragment enriched of consensus hnRNPM-binding sites (hnRNPM-BSs) within intron 45 of the NCOR2 pre-mRNA (Supplementary Fig. 8d). Indeed, hnRNPM bound to the fragment as demonstrated in RNA pulldown and RIP assays (Fig. 6a, b). Instructively, knockdown of hnRNPM enhanced the NCOR2-202 generating AS event (Fig. 6c, d), recapitulating the effect of knockdown of PLANE (Fig. 3d, e), whereas hnRNPM overexpression reduced the AS event, which was however diminished by PLANE knockdown (Fig. 6e, f), suggesting that hnRNPM is involved in PLANE-mediated regulation of NCOR2 pre-mRNA AS. In support, knockdown of PLANE reduced the amount of hnRNPM associated with the NCOR2 pre-mRNA (Fig. 6g), indicating that PLANE is necessary for the binding between hnRNPM and the pre-mRNA. In contrast, knockdown of hnRNPM did not affect the association between PLANE and the NCOR2 pre-mRNA (Fig. 6h), demonstrating that the interaction between PLANE and the pre-mRNA does not require hnRNPM. Consolidating the role of PLANE in the interaction between hnRNPM and the hnRNPM-BSs, restoration of the NCOR2-202 generating AS event by introduction of PLANE-R into cells with endogenous PLANE knockdown was associated with reinstatement of the association between hnRNPM and the hnRNPM-BSs (Figs. 4j, k & 6g). However, introduction of a PLANE mutant with the 331-751 fragment deleted to disrupt its interaction with hnRNPM or with its DFOs deleted to interfere with its interaction with the NCOR2 pre-mRNA did not restore the hnRNPM-hnRNPM-BS association (Fig. 6i). Collectively, these results indicate that PLANE facilitates the binding of hnRNPM with the NCOR2 pre-mRNA and is necessary for hnRNPM-mediated regulation of NCOR2 pre-mRNA AS.

**Figure 6.**
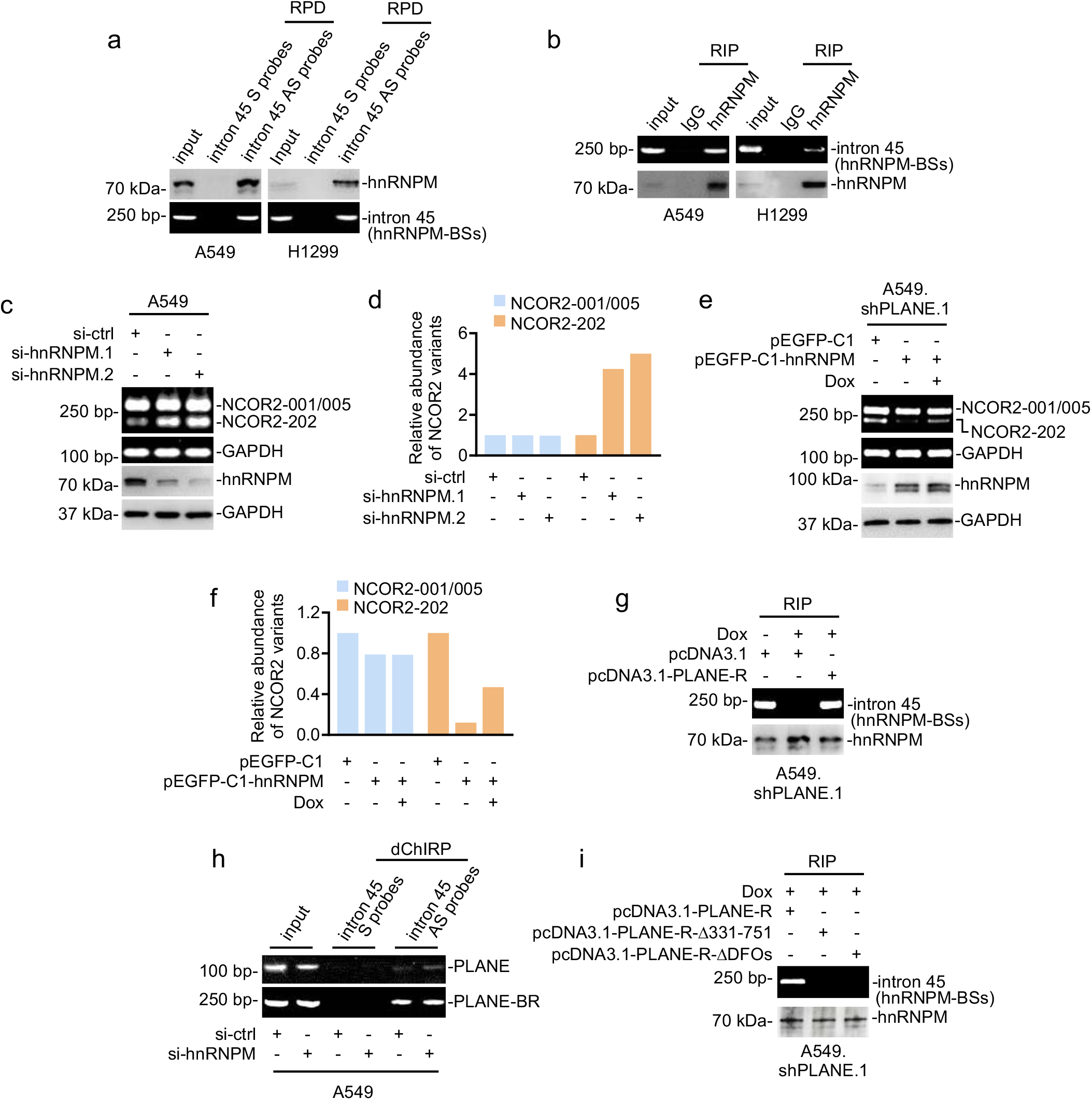
PLANE links hnRNPM to regulation of NCOR2 pre-mRNA AS. **a** hnRNPM was co-pulled down with the NCOR2 pre-mRNA using antisense probes directed to the hnRNPM binding sites (hnRNPM-BSs) at intron 45 in A549 and H1299 cells as shown in RNA pulldown assays. Data are representatives of 3 independent experiments. S, sense; AS, antisense. **b** The hnRNPM-BSs at intron 45 of the NCOR2 pre-mRNA was coprecipitated with hnRNPM in A549 and H1299 cells using RIP assays. Data are representatives of 3 independent experiments. **c** SiRNA knockdown of hnRNPM enhanced the NCOR2-202-generating AS event in A549 cells. Data are representatives of 3 independent experiments. **d** Relative levels of NCOR2-202/-001/-005 in cells with or without hnRNPM knockdown as shown in **c**. Data are representatives of 3 independent experiments. **e** Overexpression of hnRNPM reduced the NCOR2-202-generating AS event, which was reversed by co-knockdown of PLANE in A549 cells. Data shown are representatives of 3 independent experiments. **f** Relative levels of NCOR2-202/-001/-005 as shown in **e** quantitated using densitometry. Data are representatives of 3 independent experiments. **g** Induced knockdown of PLANE decreased the amount of hnRNPM associated with the hnRNPM-BSs at the NCOR2 pre-mRNA, which was reversed by co-overexpression of a shRNA-resistant PLANE mutant (PLANE-R) as shown in RIP assays. Data are representatives of 3 independent experiments. **h** PLANE was coprecipitated with a fragment at intron 45 of NCOR2 pre-mRNA containing the PLANE-BR in A549 cells with or without siRNA knockdown of hnRNPM as shown using dChIRP assays. Data are representatives of 3 independent experiments. **i** Introduction of a shRNA-resistant PLANE mutant (PLANE-R) but not a PLANE mutant with its fragment 331-751 deleted (PLANE-R-Δ331-751) or its DFOs deleted (PLANE-R-ΔDFOs) induced the association between hnRNPM and the hnRNPM-BSs at the NCOR2 pre-mRNA in A549.shPLANE cells with induced knockdown of PLANE. Data are representatives of 3 independent experiments.

## DISCUSSION

A number of proteins encoded by genes located to the distal portion of chromosome 3q that is frequently amplified in various cancer types are known to drive cancer pathogenesis, such as the p110α subunit of phosphatidylinositol 3-kinase (PI3Kα) and eukaryotic translation initiation factor 4 G (eIF4G)^44, 45^. In this study, we demonstrate that PLANE, a lncRNA encoded by a gene situated in this region, is similarly upregulated in diverse cancer types and promotes cancer cell proliferation and tumorigenicity, thus uncovering a hitherto unrecognised oncogenic contribution of a non-protein coding component of the distal portion of chromosome 3q. Nevertheless, genomic amplification is not the only mechanism responsible for the increased PLANE expression in cancer cells, rather, PLANE upregulation is more commonly driven by E2F1-mediated transcriptional activation. As a transcription factor with dichotomous functions, E2F1 on one hand transactivates many protein-coding genes involved in cell cycle progression and its high expression causes tumorigenesis^46, 47^, but on the other hand, E2F1 loss has also been demonstrated to induce cancer development and progression^48^. Our results identified transcriptional activation of PLANE as a mechanism involved in E2F1 promotion of cell proliferation, suggesting that PLANE may represent a potential target for counteracting the cancer-promoting axis of E2F1 signalling.

PLANE promoted cancer cell proliferation and tumorigenicity through inhibition of the expression of NCOR2, which, as a transcriptional corepressor, functions by way of a platform that links chromatin modifying enzymes such as HDACs and transcription factors to regulate transactivation of downstream genes involved in many cellular processes including cell survival and proliferation^9, 10, 11^. As such, deregulation of NCOR2 is associated the pathogenesis of various diseases including cancer^14, 15, 16, 17, 18^. In support of our results, a number of studies have demonstrated a tumour suppressive role of NCOR2 in cancers, such as LUAD, head and neck squamous cell carcinoma, non-Hodgkin lymphoma and osteosarcoma^14, 15, 16, 17^. However, NCOR2 has also been shown to promote cell survival and proliferation in some other cancer types such as breast cancer^18^, suggestive of an oncogenic role. These paradoxical observations are nevertheless consistent with the notion that NCOR2 functions in a manner closely related to the diverse repertoire of NCOR2 isoforms generated by AS at its C-terminal interaction domains resulting in varying affinities for different transcription factors^9, 12, 13^. Indeed, we found that PLANE-mediated suppression of NCOR2 expression was due to selective repression of AS production of the NCOR2-202 transcript that encodes one of the major NCOR2 protein isoforms, NCOR2 isoform 2^36^, demonstrating a tumour suppressive function of this isoform. Nonetheless, whether each of the other NCOR2 isoforms has any specific effect on cancer pathogenesis remains to be defined. The expression of the NCOR2 isoform BQ323636.1 is known to confer chemoresistance in breast cancer^49^. Moreover, the NCOR2 pre-mRNA splicing pattern may change in a context-dependent fashion as it does during adipocyte differentiation^50^.

Mechanistic investigations revealed that the formation of an RNA-RNA duplex was necessary for PLANE to repress the AS event generating NCOR2-202, acting through the DFOs within PLANE and the complementary PLANE-BR at intron 45 of NCOR2 pre-mRNA. The nature of the duplex interaction was substantiated through several independent experimental approaches and importantly it was demonstrated that disruption of the duplex was sufficient to abolish the repression of the NCOR2-202-generating AS event and inhibition of cell proliferation. A number of lncRNAs have been shown to regulate AS through forming RNA-RNA duplexes with pre-mRNAs and thus regulate gene expression through modification of the transcript landscape^24, 28, 51^. This is commonly accomplished through targeting particular splicing factors by lncRNA to selective cis-acting nucleotide sequences^24^. For example, the lncRNA SAF complexes with the Fas pre-mRNA at the exon 5-6 and exon 6-7 junction and binds to SPF45, thus facilitating exclusion of exon 6 leading to the generation of a Fas protein isoform that lacks the transmembrane domain and cannot induce apoptosis^28^. Moreover, the lncRNA zinc finger E-box binding homeobox 2 (ZEB2) antisense RNA 1 (ZEB2-AS1) forms RNA-RNA duplex with the ZEB2 pre-mRNA and thus prevents the recognition of the spliceosome resulting in intron inclusion at the ZEB 5’-UTR and consequent promotion of ZEB2 translation, activating EMT^51^. Similarly, the RNA-RNA duplex formed by PLANE and the NCOR2 pre-mRNA functions to facilitate the association of hnRNPM with intron 45 of the pre-mRNA leading to suppression of the NCOR2-202 generating AS event. Noticeably, PLANE was also found to regulate many other AS events, although the biological consequences remain to be defined. Similarly, the mechanisms regulating the formation of RNA-RNA duplex between PLANE and selective pre-mRNAs remain to be determined. Regardless, our results clearly demonstrated that the effect of PLANE on cancer cell proliferation is largely due to its regulation of the NCOR2-202-generating AS event through forming an RNA-RNA duplex with the NCOR2 pre-mRNA.

Like other hnRNP family proteins, hnRNPM is involved in regulating RNA maturation processes^40, 41^. In particular, it controls AS of a variety of pre-mRNAs of cancer-associated genes^40, 52, 53, 54^. For instance, hnRNPM mediates AS of fibroblast growth factor receptor 2 (FGFR2) and CD44 pre-mRNAs to promote EMT^52, 53, 54^. Moreover, hnRNPM guides an AS program involving multiple pre-mRNAs to confer resistance of Ewing sarcoma cells to inhibition of the PI3K/AKT/mTOR signal pathway^25^. Consistent with these cancer-promoting actions, our results showed that hnRNPM along with PLANE represses the AS event generating NCOR2-202 and thus promotes cancer cell proliferation. Although hnRNPs commonly act as splice suppressors when binding to exonic motifs, and as splicing enhancers when associating with motifs in introns^55, 56^, its binding to the hnRNPM-BSs at intron 45 of the NCOR2 pre-mRNA results in suppression of the NCOR2-202-generating AS event, suggesting that hnRNPM can function as an AS suppressor when it binds to intronic regulatory sequences. The finding that PLANE links hnRNPM to repress the NCOR2-202-generating AS event indicates that PLANE determines the selectivity of hnRNPM on the splice site of the NCOR2 pre-mRNA. Indeed, there is increasing evidence showing that lncRNAs can function as “local address codes” to target regulatory proteins to nucleotide sequences^37^.

The hnRNPM protein is highly abundant in cells^25, 57^. A previous quantitative proteomic study in U2OS cells showed that hnRNPM was present with >1,400,000 molecules^57^. On the other hand, PLANE is expressed at markedly lower abundance, approximately 157 to 262 molecules per A549 and H1299 cells. A question arising from this disparity is whether the limited numbers of PLANE molecules are sufficient to link enough hnRNPM necessary for repressing the NCOR2-202-generating AS event. Nevertheless, the function of hnRNPM as a splice factor is tightly regulated by its posttranslational modifications, for example, phosphorylation and sub-nuclear translocation^41^. Indeed, the vast majority of hnRNPM was found to locate to the nuclear speckles, where the pre-mRNA splicing machinery is assembled, modified and stored^42^, whereas the execution of a specific AS event is highly dynamic and conceivably requires limited resources from the general splicing machinery^42^. Thus, the factual difference between the number of PLANE copies and the number of hnRNPM molecules is conceivably not as large as estimated at the face value. Regardless, that PLANE links hnRNPM to repression of the NCOR2-202-generating AS is not in dispute since the process can be modelled *in vitro* and the action is sufficient to suppress cancer cell survival tumorigenicity.

One of the features of lncRNAs compared to protein-coding genes is their relatively poor sequence conservation^29, 37^. By use of bioinformatics analysis, we identified a Gorilla transcript that is highly homologous to human PLANE with 92% sequence similarity (Supplementary Table 5), suggesting evolutionary conservation of PLANE between Hominidae. Nonetheless, no similarity was found between PLANE and Mus musculus, a finding that precluded further testing the role of PLANE in transgenic mouse models (Supplementary Table 5). Irrespectively, our results from functional and correlative studies using human cell line models and human tissue samples suggest that PLANE contributes to cancer development and progression driven by genomic amplification of the distal portion of chromosome 3q and the cancer-promoting axis of E2F1 signalling (Supplementary Fig. 9). PLANE may therefore represent a potential anti-cancer target for counteracting these oncogenic anomalies.

## METHODS

### Cell culture and human tissues

A549, MCF-7, HCT116, Eca109, and CCC-HIE-2 cells were maintained in DMEM (Biological Industries, #01-052-1ACS; Beit Haemek, Israel) supplemented with 10% fetal bovine serum (FBS, Biological Industries, #04-001-1A; Beit Haemek, Israel) and 1% penicillin-streptomycin (Biological Industries, #03-031-1BCS, Beit Haemek, Israel). H1299, NCI-H1975 and NCI-H226 cells were cultured in RPMI-1640 (Biological Industries, #01-100-1ACS; Beit Haemek, Israel) with 10% FBS and 1% penicillin-streptomycin. CCC-HSF-1 cells were cultured in DMEM/F12 (Biological Industries, #01-172-1ACS; Beit Haemek, Israel) supplemented with 10% FBS and 1% penicillin-streptomycin. Cells were cultured in a humidified incubator at 37 °C and 5% CO_2_. All cell lines were verified to be free of mycoplasma contamination every 3 months. Individual cell line authentication was confirmed using the AmpFISTR Identifiler PCR Amplification Kit (ThermoFisher Scientific, #4427368) from Applied Biosystems and GeneMarker V1.91 software (SoftGenetics LLC). Information on cell lines is provided in Supplementary Table 6. Formalin-fixed paraffin-embedded (FFPE) normal colon mucosa, colon adenoma, COAD, LUAD and LUSC tissues were retrieved from archives of the Department of Pathology at Shanxi Cancer Hospital (Taiyuan, China). Studies using human tissues were approved by the Human Research Ethics Committees of the Shanxi Cancer Hospital in agreement with the guidelines set forth by the Declaration of Helsinki.

### Antibodies and reagents

Information on antibodies and reagents used in this study is provided in Supplementary Tables 7 & 8, respectively.

### SiRNAs and short hairpin RNA (shRNA) Oligos

SiRNAs were obtained from GenePharma (Shanghai, China) and transfected using the lipofectamine 3000 Transfection Kit (ThermoFisher Scientific, #L3000-015). ShRNA oligos were purchased from TSINGKE Biological Technology (Beijing, China).

### Plasmids

The FH1-tUTG plasmid was a kind gift from A/Professor M. J. Herold (Walter and Eliza Hall Institute of Medical Research, Australia). The pcDNA3.1(+), pGL4.73[hRluc/SV40] and pSin-3×Flag-E2F1 plasmids were kind gifts from Professor Mian Wu (Translational Research Institute, Henan Provincial People’s Hospital and People’s Hospital of Zhengzhou University, Zhengzhou, China). The pEGFP-C1 plasmid was a kind gift from A/Professor Yongyan Wu (Department of Otolaryngology, Shanxi Key Laboratory of Otorhinolaryngology Head and Neck Cancer, the first affiliated hospital, Shanxi Medical University, Taiyuan, China). The pMDLg/pRRE plasmid (#12251), pMD2.g plasmid (#12259) and pRSV-Rev plasmid (#12253) were purchased from Addgene. The pGL3-*PLANE*-promoter and the pGL3-*PLANE*-promoter-ΔE2F1-BR were purchased from Sangon Biotech (Shanghai, China). Other plasmids used in this study were generated by inserting the PCR products to the pcDNA3.1(+) or pEGFP-C1 vectors. Primers used in the fusion PCR are shown in Supplementary Table 9.

### Quantitative PCR (qPCR)

Total RNA was extracted from cultured cells using the Gene JET RNA Purification Kit (ThermoFisher Scientific, #K0731) according to the manufacturer’s instructions. cDNA was synthesized from 1 μg of total RNA using the PrimScriptTM RT reagent Kit with gDNA Eraser (TaKaRa, #RR047A; Dalian, China). Of the resultant cDNA, 12.5 ng was used in the 20 μl qPCR mix, containing 10 μl of TB Green Premix Ex Taq II (Tli RNaseH Plus) (TaKaRa, #RP820A; Dalian, China) and 0.4 μM of each primer. Samples were amplified for 40 cycles using a StepOnePlus™ Real-Time PCR System (ThermoFisher Scientific). 2^−ΔΔCT^ method was used to calculate the relative gene expression levels normalized to the GAPDH housekeeping control. Primer sequences are listed in Supplementary Table 10.

### Chromatin Immunoprecipitation (ChIP)

ChIP assays were performed using the ChIP Assay Kit (Beyotime, #P2078; Shanghai, China) according to the manufacturer’s instructions. Briefly, cells were cross-linked with a final concentration of 1% formaldehyde in growth medium for 15 min at 37 °C and quenched by the addition of glycine solution for 5 min at room temperature (RT). Then cells were harvested, lysed and sonicated. After being cleared by centrifugation at 12,000 × g for 10 min at 4 °C, the cell lysate was subjected to a 1:10 dilution and rotated with E2F1, H3K4me3 and H3K27me3 antibodies or corresponding mouse/rabbit normal immunoglobulin (IgG) antibodies at 4 °C overnight. Then, 60 μl of protein A/G agarose beads was added to the antibody-lysate mixture and rotated at 4 °C for an additional 1 hr. Beads were washed, and DNA fragments were eluted, purified and subjected to PCR analysis using the specific primers. PCR products were separated by gel electrophoresis on the 2% agarose gel. Information on antibodies and primers used in this study are shown in Supplementary Tables 7 & 11, respectively.

### Luciferase reporter assays

A549 and H1299 cells were transfected with pGL3-*PLANE*-promoter reporters or pGL3-*PLANE*-promoter-ΔE2F1-BR reporters together with pGL4.73[hRluc/SV40] reporters expressing the renilla luciferase. After 48 hr, firefly and renilla luciferase activities were examined by a Dual-Luciferase® Reporter Assay System (Promega, #E910) with a VARIOSKAN LUX microplate reader. The renilla luciferase activity was used to normalize the firefly luciferase activity.

### Colony formation

Cancer cells were seeded in six-well plates at 2,000 cells/well. After growing for further two weeks, cells were fixed with methanol and staining with 0.5% crystal violet. The images were captured with a Bio-Rad GelDoc™ XR + imaging system (Bio-Rad). The percentage and intensity of area covered by crystal violet-stained cell colonies were quantified using ImageJ-plugin “ColonyArea”.

### Cell cycle analysis

Cell cycle analysis was performed using the Cell Cycle and Apoptosis Analysis Kit (Meilunbio, #MA0334; Dalian, China) according to the manufacturer’s instructions followed by flow cytometry. Briefly, A549 and H1299 cells transfected with PLANE siRNAs for 48 hr in 24-well plates were harvested and fixed in 75% ethanol at 4 °C overnight. After being centrifuged, cells were incubated in the staining solution at 37 °C in the dark for 30 min. Then cells were subjected to analysis using a flow cytometer (FACSAria, BD Biosciences).

### Anchorage-independent cell growth

Cells carrying an inducible PLANE knockdown in response to Dox were seeded in the Ultra-Low attachment 6-well plate (Corning, #3471) at 2,000 cells/well. Cells with or without treatment with doxycycline (Dox) and cessation of Dox treatment were incubated at 37 °C in a humidified incubator until colonies were formed. Colonies were counted under a light microscope^29^.

### In situ hybridization (ISH)

ISH assays were performed using the RNAscope^®^ 2.5 HD Detection Reagent-BROWN (Advanced Cell Diagnostics, #322310) according to the manufacturer’s instructions^37, 58^. Briefly, FFPE LUSC and LUAD as well as COAD tissue microarrays (#HLug-Squ150Sur-02, #HLugA180Su03, #HColA180Su12) purchased from the Shanghai Outdo Biotech Co., Ltd (China) were deparaffinized in xylene for 5 min at RT twice, followed by dehybridization in 100% alcohol. After being air-dried, the tissue sections were incubated with hydrogen peroxide for 10 min at RT and washed in the distilled water for five times. Then the sections were heated in target retrieval reagent to 100 °C for 20 min, followed by being treated with proteinase K and incubated in hybridization buffer containing probes (Advanced Cell Diagnostics, #570031) at 40 °C for 3 hr. After being washed, the sections were incubated with 3,3’-diaminobenzidine (DAB), and counterstaining was carried out using hematoxylin.

The percentage of positive cells was ranged from 0 to 100%. The intensity of staining (intensity score) was judged on an arbitrary scale of 0 to 4: no staining (0), weakly positive staining (1), moderately positive staining (2), strongly positive staining (3) and very strong positive staining (4). A reactive score (RS) was derived by multiplying the percentage of positive cells with staining intensity divided by 10.

### Immunofluorescence (IF)

Cells grown on coverslips were fixed in 4% formaldehyde for 10 min at RT. After being washed using PBS, cells were then permeabilized in blocking buffer for 60 min at RT. Antibodies diluted 1:500 in blocking buffer were incubated with cells overnight at 4 °C. Cells were washed in PBS and incubated with secondary antibodies diluted 1:200 in blocking buffer for 60 min at RT in the dark. After being washed, cells were mounted in the ProLong™ Glass Antifade Mountant with NucBlue reagent (ThermoFisher Scientific, P36981). Images were digitally recorded using a Leica SP8 confocal microscope. Information of antibodies used in this study was shown in Supplementary Table 7.

### Subcellular fractionation

Cells were harvested by trypsinization and lysed in hypotonic buffer A (10 mM Hepes pH 7.9, 10 mM KCl, 0.1 mM EDTA, 0.1 mM EGTA, 1 mM DTT, 0.15% Triton X-100, cOmplete™, EDTA-free Protease Inhibitor Cocktail) on ice for 15 min. The supernatants after centrifugation at 12,000 × g for 3 min were collected as the cytoplasmic fractions and the pellets were subjected to the nuclear fractionation. The pellets were rinsed with cold PBS once and lysed in an equal volume of buffer B (20 mM Hepes pH 7.9, 400 mM NaCl, 1 mM EDTA, 1 mM EGTA, 1 mM DTT, 0.5% Triton X-100, cOmplete™, EDTA-free Protease Inhibitor Cocktail) on ice for 15 min. Cytoplasmic and nuclear fractions were centrifuged at 16,000 × g for 20 min to remove the insoluble debris. The supernatants were collected for RNA isolation and immunoblotting analysis.

### *In vitro* transcription

The DNA templates used for *in vitro* synthesis of PLANE, antisense PLANE and NCOR2 pre-mRNA were generated by PCR amplification from cDNAs using PrimerSTAR Max DNA Polymerase (TAKARA, #R045A; Dalian, China). Forward primers containing the T7 RNA polymerase promoter sequence and reverse primers without the promoter sequence were used for synthesizing PLANE, antisense PLANE, and NCOR2 pre-mRNA. After PCR amplification, the products were purified using a MiniBEST Agarose Gel DNA Extraction Kit (Takara, #9762; Dalian, China), and subjected to *in vitro* transcription using a TranscriptAid T7 High Yield Transcription Kit (ThermoFisher Scientific, #K0441) according to the manufacturer’s instructions. The *in vitro-*transcribed RNAs could be further labelled with biotin using a Pierce™ RNA 3’ End Desthiobiotinylation Kit (ThermoFisher Scientific, #20163). Primer sequences are shown in Supplementary Table 11.

### Domain-specific chromatin isolation by RNA purification (dChIRP)

dChIRP assays were performed as previous described^59^. Briefly, A549 and H1299 cells were harvested and cross-linked in 1% glutaraldehyde for 10 min at RT with rotation. The cross-linked cells were lysed in lysis buffer (50 mM Tris-Cl [pH 7.0], 10 mM EDTA, 1% SDS, PMSF, Superase-in), followed by sonication. 4 μg antisense / sense biotin-labelled probes or 10 μg *in vitro-*transcribed biotin-labelled PLANE / antisense PLANE were rotated with cell lysates at 37 °C for 4 hr, followed by adding 100 μl C-1 magnetic beads (Invitrogen, #65002) to each sample and incubating at 37 °C for 30 min with rotation. Beads were then washed in wash buffer for five times, followed by RNA isolation. Probe sequences are shown in Supplementary Table 11.

### Biotin RNA pull-down (RPD)

A549 and H1299 cells were harvested and washed in PBS for three times. Cell pellets were then lysed in lysis buffer (50 mM Tris-HCl [pH 7.5], 150 mM NaCl, 2.5 mM MgCl_2_, 1 mM EDTA, 10% Glycerol, 0.5% Nonidet P-40/Igepal CA-630, 1 mM DTT, cOmplete™ EDTA-free Protease Inhibitor Cocktail and RNase inhibitors) and sonicated. 4 μg antisense / sense biotin-labelled probes were incubated with lysates at 4 °C overnight before rotating with streptavidin beads (ThermoFisher Scientific, #20349) for additional 2 hr. Beads were then washed in lysis buffer for four times, followed by RNA isolation and immunoblotting analysis. Information of antibodies and probes is shown in Supplementary Tables 7 & 11, respectively.

### Mass spectrometry (MS) analysis

Proteins co-pulled down with RNA using antisense / sense biotin-labelled probes were separated by 10% acrylamide gels and visualized by Coomassie brilliant blue staining. The specific protein band shown in the group using antisense probes along with the corresponding region in the group using sense probes were resected and digested, followed by the liquid chromatography–mass spectrometry (LS-MS) analysis using a mass spectrometer (ThermoFisher Scientific, EASY-nLC1000 & LTQ Orbitrap Velos Pro). Proteins identified from the mass spectrometry analysis are listed in Supplementary Table 4.

### RNA immunoprecipitation (RIP)

RIP assays were performed using a Magna RIP™ Kit (Millipore, #17-700; Darmstadt, Germany) according to the instruction provided by the manufacturer. Briefly, cell lysates prepared in hypotonic buffer supplemented with RNase inhibitor and protease inhibitor were incubated with magnetic beads pre-incubated with hnRNPM antibodies at 4 °C overnight. After being washed with RIP wash buffer, the bead-bound immunocomplexes were subjected to immunoblotting analysis and RNA isolation. Information on antibody and primers used in this study is shown in Supplementary Tables 7 & 11.

### Immunoprecipitation (IP)

Cells were collected with trypsinization and lysed with lysis buffer (20 mM Tris-HCl pH 8.6, 100 mM NaCl, 20 mM KCl, 1.5 mM MgCl2, 0.5% NP-40, cOmplete™ EDTA-free Protease Inhibitor Cocktail) on ice for 1 hr and centrifuged at 16,000 × g for 30 min. After quantification using a BCA protein assay kit (ThermoFisher, #23225), 3 mg of total protein were rotated with antibodies at 4 °C overnight. Protein-antibody complexes were then captured with the Pierce™ Protein A/G Agarose (ThermoFisher Scientific, #20421) at 4°C for 2 hrs with rotation and beads were then rinsed with wash buffer (25 mM Tris, 150 mM NaCl, pH 7.2), boiled and subjected to immunoblotting analysis. Antibodies used in this study are shown in Supplementary Table 7.

### Absolute quantification of PLANE

Absolute RNA quantification was performed using the standard curve method by qPCR. cDNA was synthesized using 1 μg of the total RNA extracted from a fixed cell number. Ten-fold serial dilutions of the pcDNA3.1-PLANE plasmid (10^2^ to 10^7^ molecules per ml) were used as a reference molecule for the standard curve calculation. Assays were reconstituted to a final volume of 20 μl using 5 μl cDNA from cells or 5 μl serial diluted pcDNA3.1-PLANE plasmid and cycled using a StepOnePlus™ Real-Time PCR System. Data calculated as copies per 5 μl cDNA were converted to copies per cell based on the known input cell equivalents. Primer sequences used are listed in Supplementary Table 10.

### Inducible shRNA knockdown

The FH1-tUTG inducible knockdown vector was digested using BsmBII (NEW ENGLAND BioLabs, #R0580S) and XhoI (NEW ENGLAND BioLabs, #R0146S) enzymes, and the annealed shRNA oligos were inserted into the digested vector using the T4 DNA ligase (ThermoFisher Scientific, #EL0014). The lentiviral particles were packaged via co-transfection of FH1-tUTG vector inserted with shRNA oligos (44 μg), pMDLg/pRRE plasmid (22 μg), pMD2.g plasmid (13.2 μg) and pRSV-Rev plasmid (11 μg) plasmids into HEK293T cells^60^. A549 or H1299 cells were transduced with the lentiviral particles in 6 cm cell culture dishes to establish inducible knockdown cell sublines. The knockdown of PLANE was induced in response to doxycycline treatment. ShRNA sequences are shown in Supplementary Table 12.

### Xenograft mouse model

A549 cells expressing the inducible PLANE shRNAs were subcutaneously injected into the dorsal flanks of 4-week-old female nude mice (6 mice per group, Shanghai SLAC Laboratory Animal Co. Ltd., China). Tumor growth was measured every 3 days using a calliper. Mice were sacrificed after 33 days of cancer cell transplantation. Tumors were excised and measured. Studies on animals were conducted in accordance with relevant guidelines and regulations and were approved by the Animal Research Ethics Committee of the first affiliated hospital, Shanxi Medical University and Shanxi Cancer Hospital and Institute (China). All mice were housed in a temperature-controlled room (21-23 °C) with 40–60% humidity and a light/dark cycle of 12 h/12 h.

### Statistical Analysis

Statistical analysis was carried out using the GraphPad Prism 8 to assess differences between experimental groups. Statistical differences were analyzed by two-tailed Student’s *t*-test or one-way ANOVA test followed by Tukey’s multiple comparisons. *P* values lower than 0.05 were considered to be statistically significant.

## DATA AVAILABILITY

The RNA sequencing data have been deposited in the NCBI Gene Expression Omnibus database under the accession code GSE162215. The mass spectrometry proteomics data have been deposited to the ProteomeXchange Consortium (http://proteomecentral.proteomexchange.org) via the iProX partner repository with the dataset identifier PXD022747. The long noncoding RNA expression data and E2F1 mRNA expression data referenced during the study are available in a public repository from the Cancer RNA-seq Nexus dataset (http://syslab4.nchu.edu.tw/). The cancer patient survival data referenced during the study are available in a public repository from the GEPIA website (http://gepia.cancer-pku.cn/) under the accession codes TCGA-LUSC, TCGA-COAD, TCGA-KIRC and TCGA-UCEC. The gene amplification frequency data referenced during the study are available in a public repository from the cBioPortal website (https://www.cbioportal.org/) under the accession code TCGA PanCancer Atlas Studies.

## ACKNOWLEDGEMENTS

This work was supported by the National Health and Medical Research Council (NHMRC; APP1147271, APP1162753, APP1177087) and Cancer Council NSW Project Grant (RG20-10), Australia. The authors thank A/Professor M. J. Herold (Walter and Eliza Hall Institute of Medical Research, Australia) for the plasmid FH1tUTG, Professor Xiao Ying Liu (Translational Research Institute, Henan Provincial People’s Hospital, China) and Professor Xiaoju Zhang (Respiration Department, Henan Provincial People’s Hospital, Zhengzhou, China) for providing the Eca109 and NCI-H1975 as well as NCI-H226 cell lines, respectively. The authors thank the Shanghai luming biological technology co., LTD (Shanghai, China) for providing proteomics services.

## AUTHOR CONTRIBUTIONS

X.D.Z., F.-M.S., L.J., and T. Liu designed the experiments. X.D.Z., F.-M.S., and L.J. supervised the work. L.T., Y.C.F., P.L.W., S.X.W., T.F.Q., S.N.Z., T.La, Y.Y.Z., X.H.Z., D.Z., and J.Y.W. performed experiments using human cell lines and tissues and related data collections; S.T.G. conducted experiments in xenograft models; T.Liu, J.M.L., Y.C.F., L.J., T.La, R.F.T., and J.Y.W. carried out analysis of publicly available data and bioinformatics analysis. X.D.Z., R.F.T., F.-M.S., T.Liu, and L.J. wrote the manuscript. All authors commented on the manuscript.

