## Supplementary Figures 1-9 for "The pan-cancer lncRNA PLANE regulates an alternative splicing program to promote cancer pathogenesis"

Supplementary Figure 1

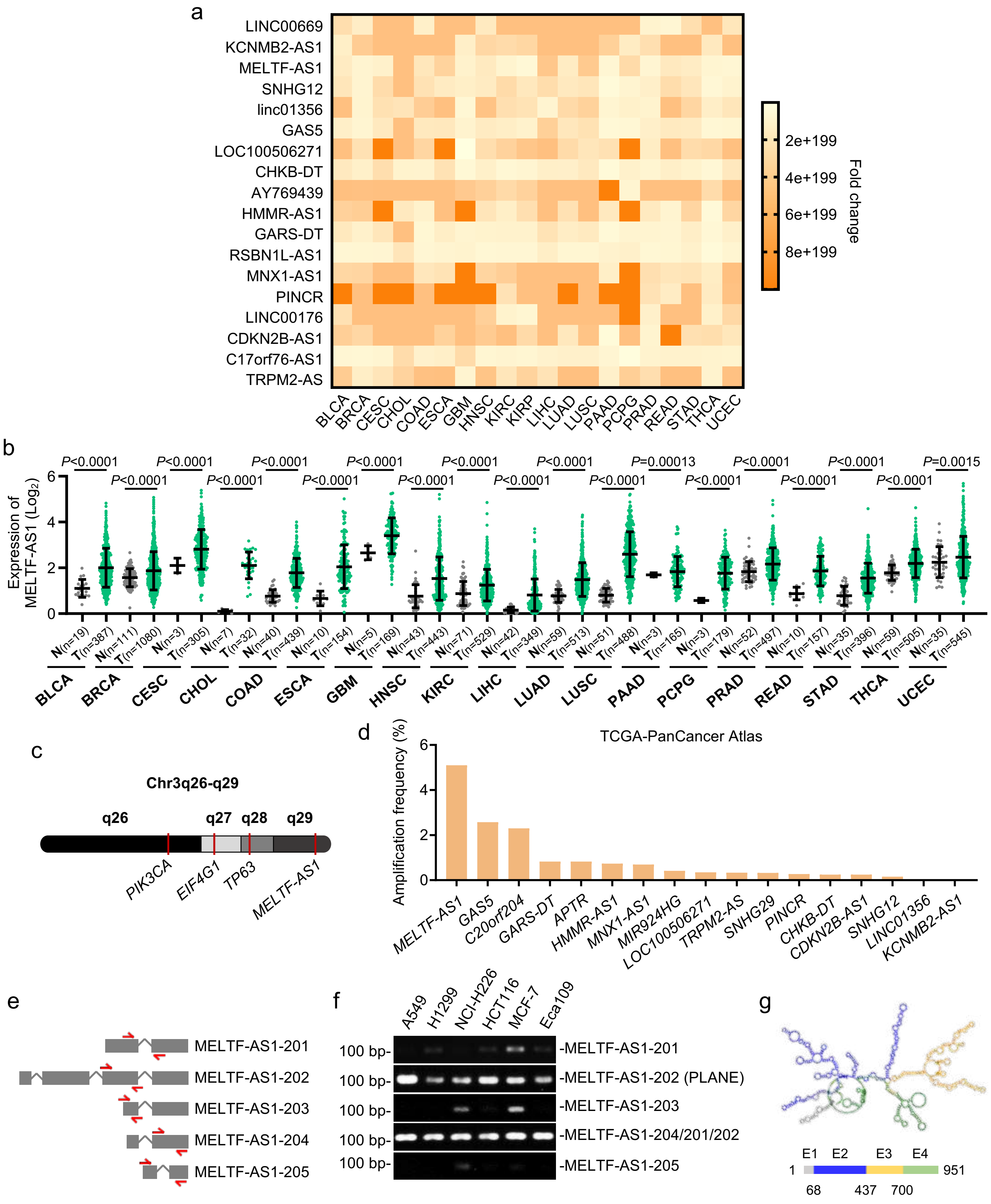

**Supplementary Figure 1. MELTF-AS1 (PLANE) is a pan-cancer-associated lncRNA**

**a** Identification of pan-cancer-associated lncRNAs through analysis of the lncRNA expression data from the TCGA dataset. Eighteen lncRNAs that are increased in at least 19 of 20 types of cancer relative to corresponding normal tissues are depicted using a heatmap. The data are fold changes in cancer compared with normal tissues. BLCA: bladder urothelial carcinoma; BRCA: breast invasive carcinoma; CESC: cervical squamous cell carcinoma and endocervical adenocarcinoma; CHOL: cholangiocarcinoma; COAD: colon adenocarcinoma; ESCA: esophageal carcinoma; GBM: glioblastoma multiforme; HNSC: head and neck squamous cell carcinoma; KIRC: kidney renal clear cell carcinoma; KIRP: kidney renal papillary cell carcinoma; LIHC: liver hepatocellular carcinoma; LUAD: lung adenocarcinoma; LUSC: lung squamous cell carcinoma; PAAD: pancreatic adenocarcinoma; PCPG: pheochromocytoma and paraganglioma; PRAD: prostate adenocarcinoma; READ: rectum adenocarcinoma; STAD: stomach adenocarcinoma; THCA: thyroid carcinoma; UCEC: uterine corpus endometrial carcinoma.

**b** MELTF-AS1 is upregulated in diverse types of cancer compared with corresponding normal tissues as revealed by analysis of the lncRNA expression data in the TCGA dataset. Data are mean ± s.d.; two-tailed Student's *t*-test. N: normal tissues; T: tumour tissues.

**c** Schematic illustration of the genomic location of the *MELTF-AS1* gene and representative protein-coding genes that are involved in cancer pathogenesis at chromosome 3q26-29.

**d** Analysis copy-number alteration data of the PanCancer Atlas of the TCGA showing that *MELTF-AS1* was the most frequently amplified gene among those that encode the pan-cancer upregulated lncRNAs as shown in **a**.

**e** Schematic illustration of the five annotated MELTF-AS1 transcript isoforms. Filled boxes represent exons. The primers used for specific detection of each isoform were depicted in red. Primers directed against MELTF-AS1-204 span a common region shared by MELTF-AS1-201 and MELTF-AS1-202.

**f** PCR analysis showing that the longest isoform of MELTF-AS1, MELTF-AS1-202 (PLANE) was markedly more abundant than other isoforms in the indicated cancer cell lines. Data are representatives of 3 independent experiments.

**g** The secondary structure model of PLANE is predicted based on minimum free energy algorithm.

Supplementary Figure 2

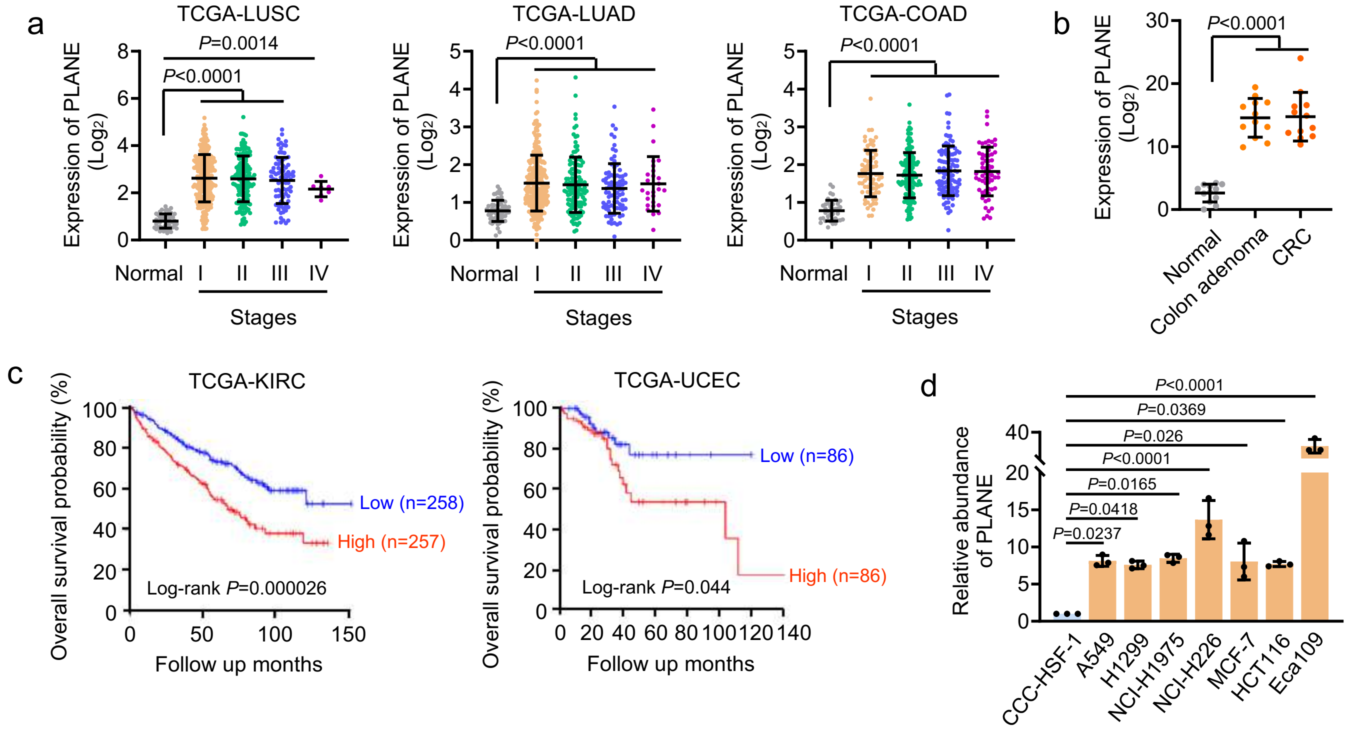


**Supplementary Figure 2. PLANE is upregulated at early stages during cancer pathogenesis**

**a** Analysis of the TCGA data showing that PLANE expression levels did not differ among tumours of different stages in LUSC, LUAD and COAD. Data are mean ± s.d.; one-way ANOVA followed by Tukey's multiple comparisons test.

**b** PLNAE expression was upregulated in colon adenoma tissues (n = 12) compared with normal colon epithelial tissues (n = 12), whereas there was no significant difference in PLANE expression levels between colon adenoma and colon cancer tissues (n = 12) as determined by qPCR. Data are mean ± s.d.; one-way ANOVA followed by Tukey's multiple comparisons test.

**c** Kaplan-Meier analysis of the probability of overall survival of kidney clear cell carcinoma (KIRC; n=515) and uterine corpus endometrial carcinoma (UCEC; n=172) patients derived from the TCGA dataset using the median of PLANE levels as the cutoff.

**d** PLANE was expressed at higher levels in the indicated cancer cell lines than the normal human fibroblast cell line CCC-HSF-1. Data are mean ± s.d.; n = 3 independent experiments, one-way ANOVA followed by Tukey's multiple comparisons test.

Supplementary Figure 3

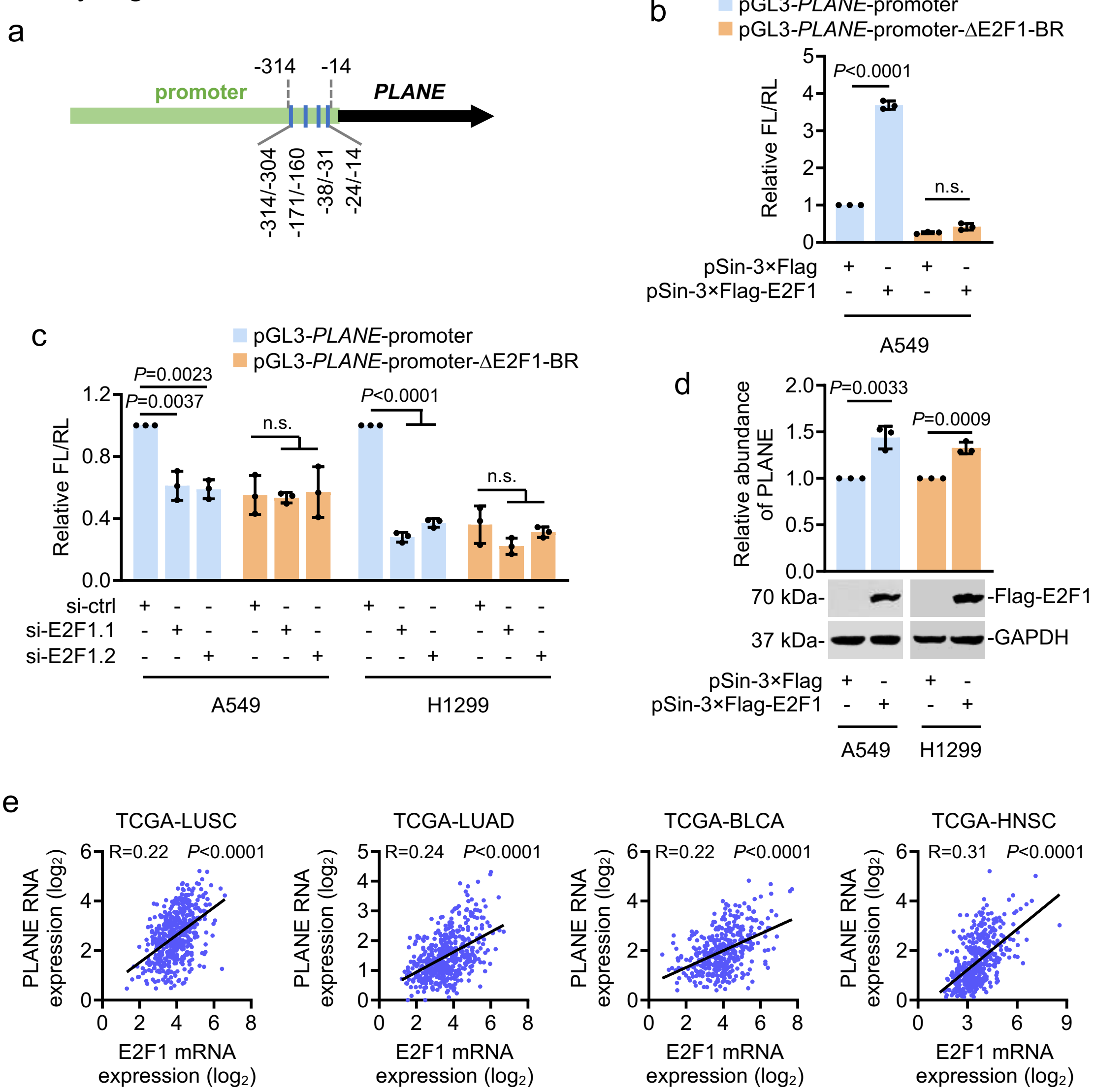


**Supplementary Figure 3. E2F1 transcriptionally activates PLANE expression**

**a** Schematic illustration of multiple consensus E2F1 binding motifs (blue bars) located to the -314/-14 region of the proximal promoter of *PLANE* gene.

**b** Overexpression of E2F1 enhanced the transcriptional activity of a *PLANE* reporter construct with the intact E2F1 binding region (BR)(pGL3-*PLANE*-promoter) but did not affect the activity of a construct with the E2F1-BR deleted (pGL3-*PLANE*-promoter-ΔE2F1-BR) in A549 cells. Data are mean ± s.d.; n = 3 independent experiments, one-way ANOVA followed by Tukey's multiple comparisons test. FL: Firefly luciferase activity; RL: Renilla luciferase activity.

**c** SiRNA knockdown of E2F1 reduced the transcriptional activity of a *PLANE* reporter construct with the intact E2F1-BR (pGL3-*PLANE*-promoter) but did not affect the activity of a construct with the E2F1-BR deleted (pGL3-*PLANE*-promoter-ΔE2F1-BR). Data are mean ± s.d.; n = 3 independent experiments, one-way ANOVA followed by Tukey's multiple comparisons test.

**d** Overexpression of E2F1 caused upregulation of PLANE in A549 and H1299 cells. Data are mean ± s.d. or representatives; n = 3 independent experiments, two-tailed Student's *t*-test.

**e** Linear regression analysis of the relationship between PLANE and E2F1 mRNA expression in the LUSC, LUAD, bladder urothelial carcinoma (BLCA) and head and neck squamous cell carcinoma (HNSC) datasets derived from the TCGA.

Supplementary Figure 4

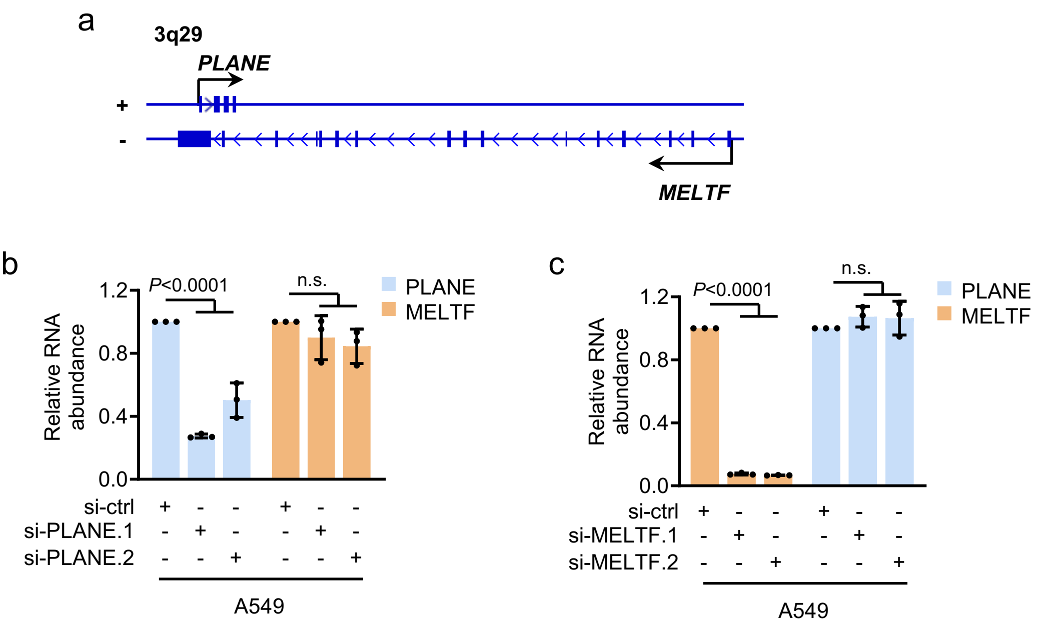


**Supplementary Figure 4. PLANE does not affect the expression of its neighbour gene *MELTF***
**a** Schematic illustration of the genomic location of the *PLANE* and *MELTF* genes.
**b** Knockdown of PLANE did not impinge on MELTF expression in A549 cells. Data are mean ± s.d.; n = 3 independent experiments, one-way ANOVA followed by Tukey's multiple comparisons test.
**c** Knockdown of MELTF did not impact PLANE expression in A549 cells. Data are mean ± s.d.; n = 3 independent experiments, one-way ANOVA followed by Tukey's multiple comparisons test.

Supplementary Figure 5

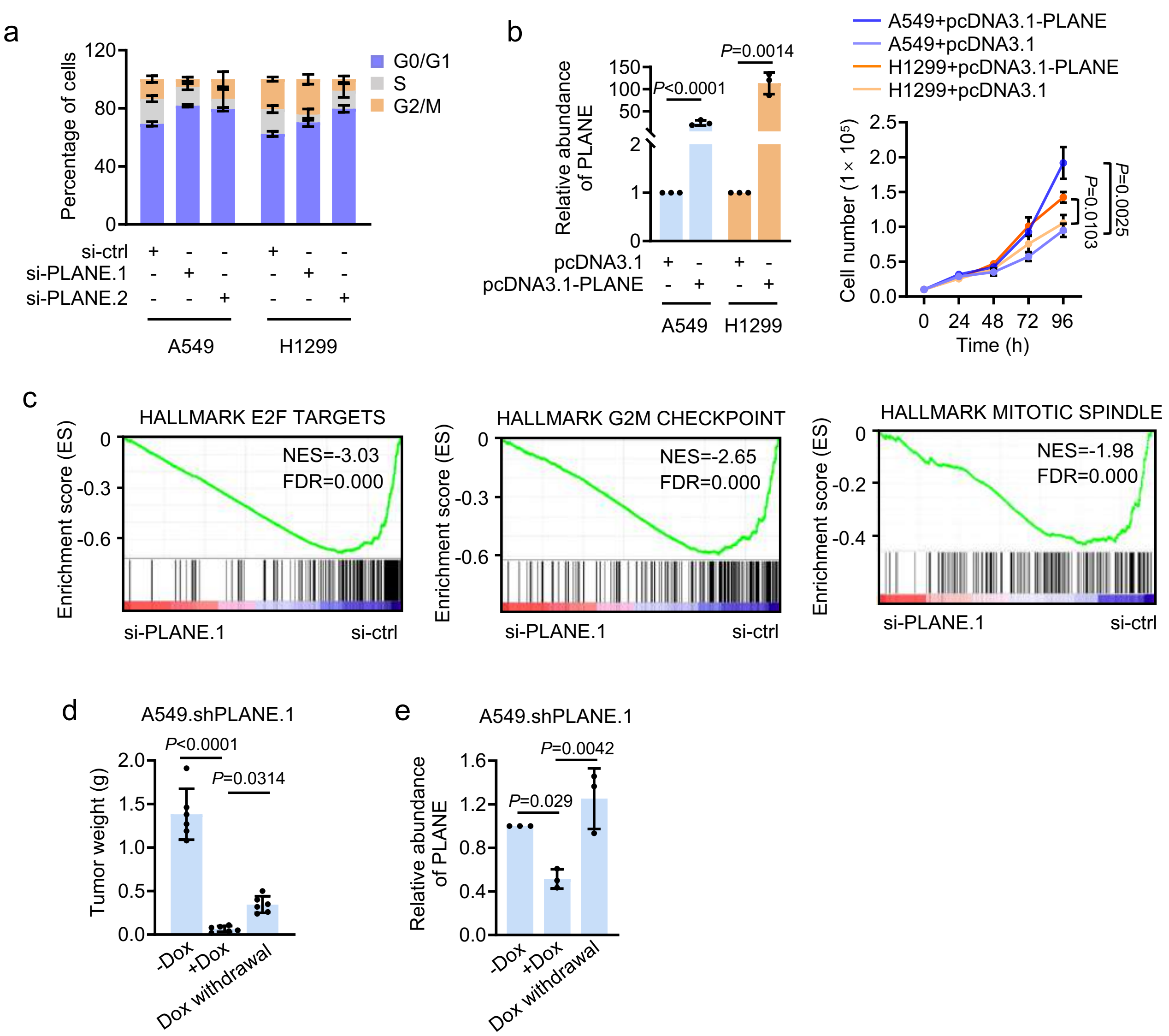


**Supplementary Figure 5. PLANE promotes cancer cell proliferation**

**a** SiRNA knockdown of PLANE caused cell cycle arrest at G0/G1 phase in A549 and H1299 cells as shown by propidium iodide staining followed by flow cytometry analysis. Data are mean ± s.d.; n = 3 independent experiments.

**b** Overexpression of PLANE (left) promoted cell proliferation as shown by accelerated cell number increases (right) in A549 and H1299 cells. Data are mean ± s.d.; n = 3 independent experiments, two-tailed Student's *t*-test.

**c** GSEA of RNA-seq data from A549 cells. n=2 experimental repeats. NES, normalised enrichment score; FDR, false discovery rate.

**d & e** Tumour weights (d) (n = 6 mice per group) and PLANE expression in representative samples (e) of A549.shPLANE xenografts in nu/nu mice with or without treatment with doxycycline (Dox, 2 mg/ml supplemented with 10 mg/ml sucrose in drinking water) and cessation of Dox treatment. Data are mean ± s.d.; one-way ANOVA followed by Tukey's multiple comparisons test.

Supplementary Figure 6

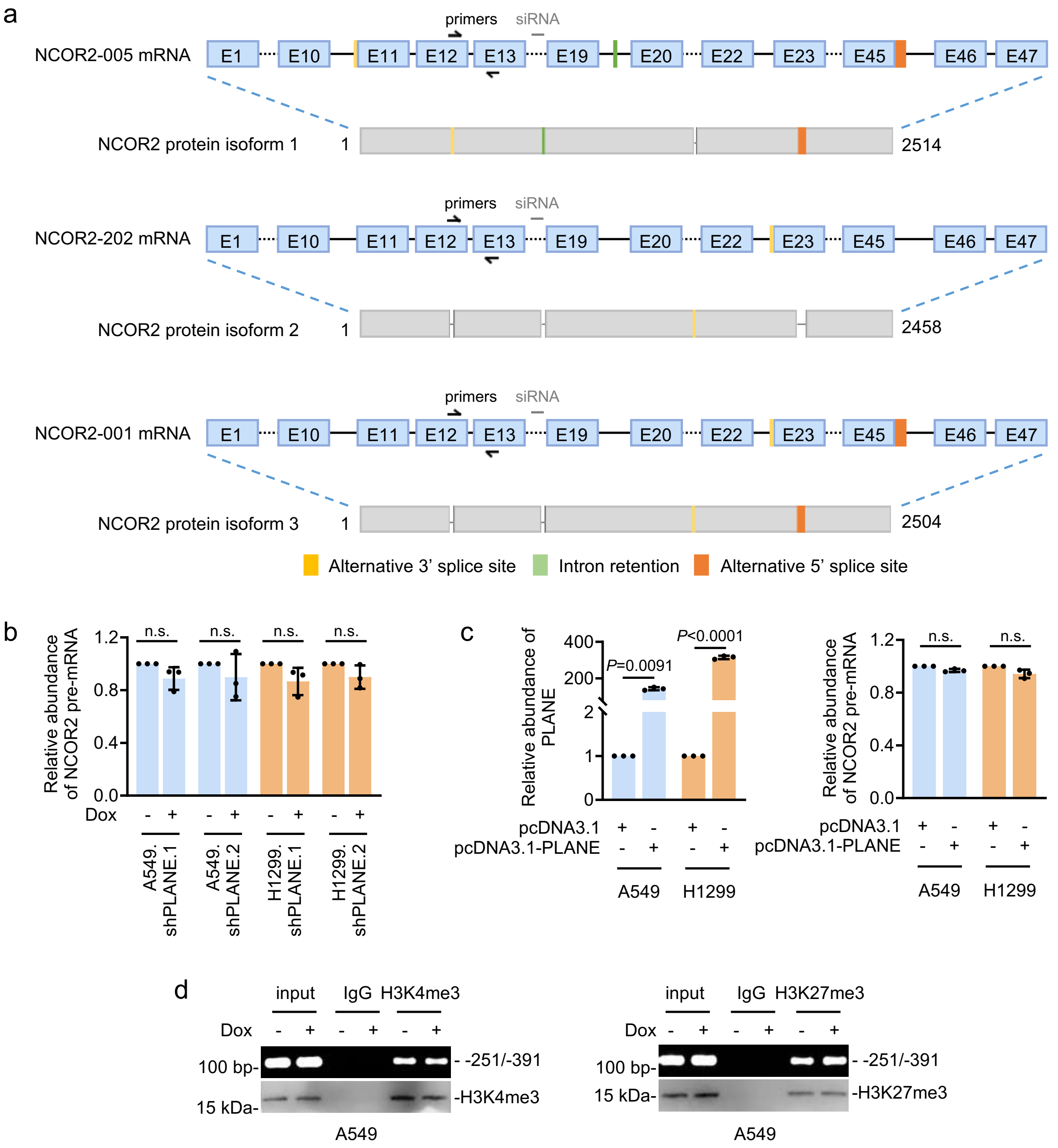


**Supplementary Figure 6. PLANE does not regulate NCOR2 transcription**

**a**, Schematic illustration of the NCOR2 protein isoforms, NCOR2 isoform 1, NCOR2 isoform 2 and NCOR2 isoform 2 that are respectively encoded by the NCOR2 mRNA variants NCOR2-005, NCOR2-202 and NCOR2-001. E: exon.

**b & c** Neither induced knockdown (b) nor overexpression (c) of PLNAE impinged on NCOR2 pre-mRNA expression. Data are mean ± s.d.; n = 3 independent experiments, two-tailed Student's *t*-test.

**d** SiRNA knockdown of PLANE did not affect the enrichment of the transcriptional activation mark H3K4me3 and the transcriptional repression mark H3K27me3 to the *NCOR2* promoter in A549 and H1299 cells as shown in chromatin immunoprecipitation (ChIP) assays. Data are representatives of 3 independent experiments.

Supplementary Figure 7

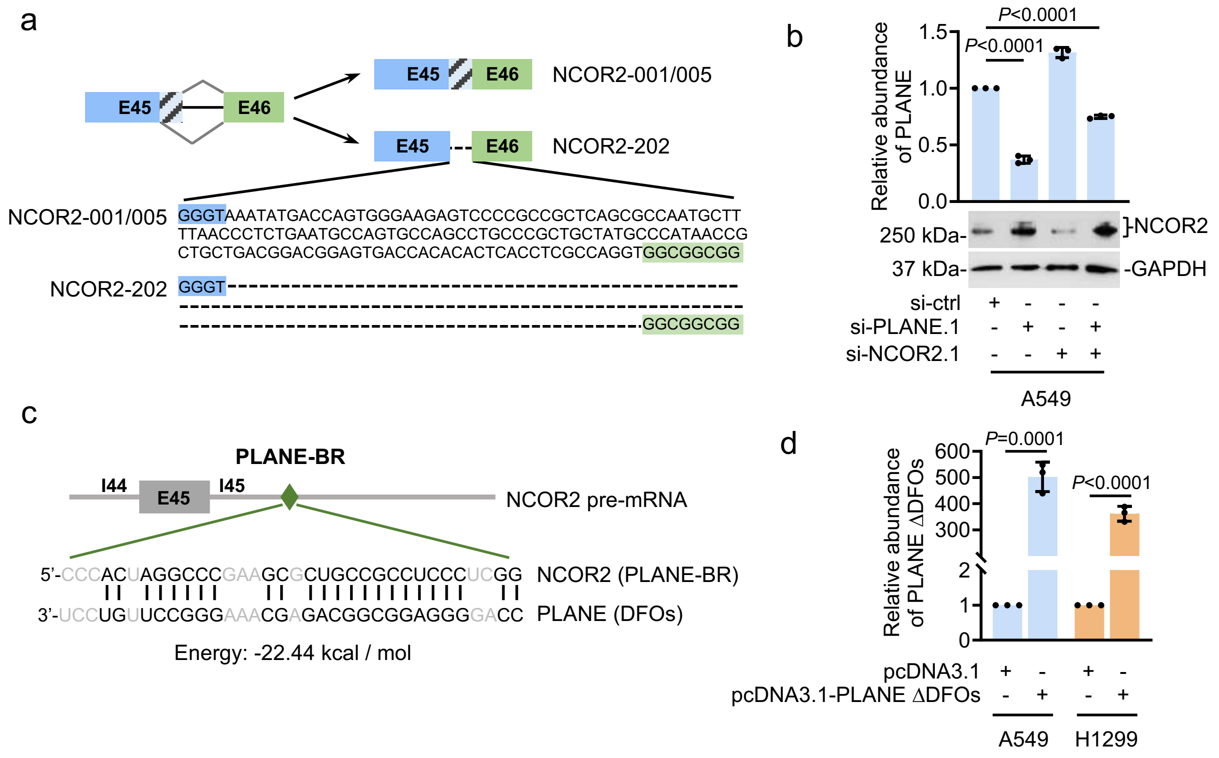


**Supplementary Figure 7. PLANE regulates NCOR2-202 pre-mRNA AS**

**a** Schematic illustration of the sequence overlaps between NCOR2-202 and NCOR2-001/-005 and the generation of NCOR2-202 through an alternative 5' splice site.

**b** SiRNA knockdown of PLANE and NCOR2 in A549 cells. Data are mean ± s.d. or representatives; n = 3 independent experiments, one-way ANOVA followed by Tukey's multiple comparisons test.

**c** Schematic illustration of base-pairing between the PLANE binding region (PLANE-BR) at intron 45 (I45) of the NCOR2 pre-mRNA and the duplex-forming oligonucleotides (DFOs) of PLANE.

**d** Overexpression of a PLANE mutant with the DFOs deleted (PLANE-ΔDFOs) in A549 and H1299 cells. Data are mean ± s.d.; n=3 independent experiments, two-tailed Student's *t*-test.

Supplementary Figure 8

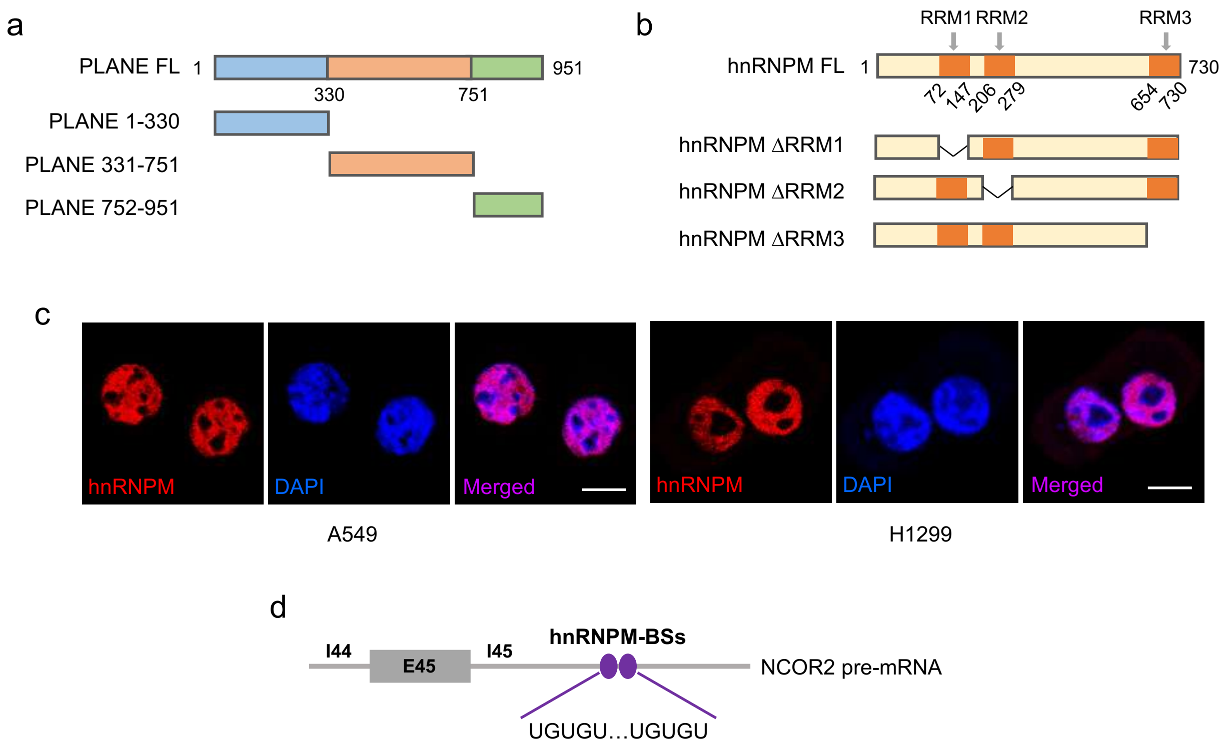


**Supplementary Figure 8. PLANE binds to hnRNPM and facilitates repression of the NCOR2-202-generating AS event by PLANE**

**a** Schematic illustration of full-length PLANE (PLANE-FL) and the PLANE mutants (PLANE 1-330, PLANE 331-751 and PLANE 752-951) used in mapping experiments.

**b** Schematic illustration of full-length hnRNPM (hnRNPM FL) and the hnRNPM mutants with individual RNA recognition motifs (RRM1, RRM2 and RRM3) deleted.

**c** Representative microscopic photographs of immunofluorescence staining of hnRNPM in A549 cells grown on coverslips. Data shown are representatives of 3 independent experiments. Scale bar: 10 µm.

**d** Schematic illustration of the consensus hnRNPM-binding sites (hnRNPM-BSs) at intron 45 of the NCOR2 pre-mRNA.

Supplementary Figure 9

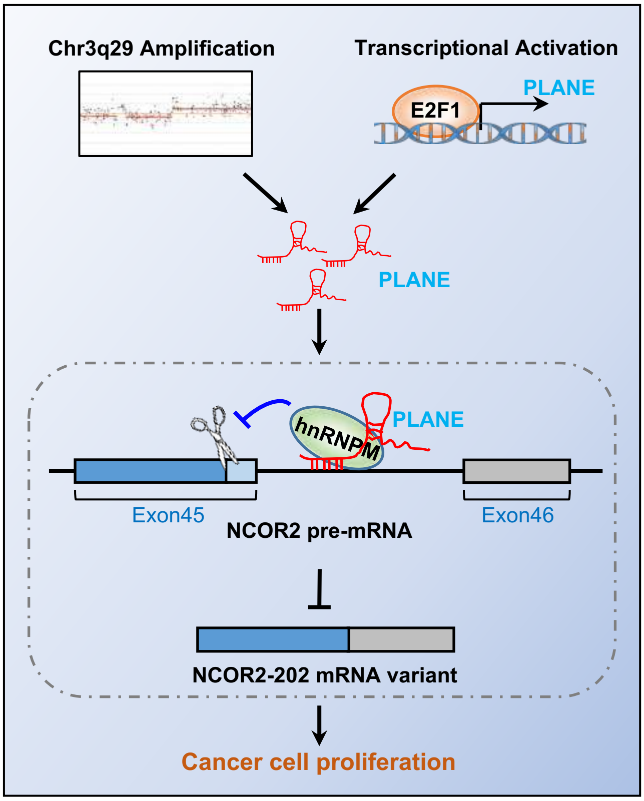


**Supplementary Figure 9. Schematic illustration PLANE facilitation of hnRNPM-mediated repression of the NCOR2-202-generating alternative splicing (AS) event to promote tumorigenesis**

PLANE is upregulated in diverse cancer types driven by genomic amplification and transcriptional activation by E2F1 and forms RNA-RNA duplex with the NCOR2 pre-mRNA and binds to hnRNPM. This facilitates the association of hnRNPM with the NCOR2 pre-mRNA at intron 45, leading to repression of the NCOR2-202-generating alternative splicing event, leading to promotion of tumorigenesis.
