## Supplementary Tables 1-12 for "The pan-cancer lncRNA PLANE regulates an alternative splicing program to promote cancer pathogenesis"

**Supplementary Table 1. Summary of clinicopathological characteristics of the cohort of 75 lung squamous cell carcinoma patients**

| **Characteristics** | **Cases** | **PLANE abundance in Lung squamous cell carcinoma(RS^1^)** | | ***P* value^2^** |
| --- | --- | --- | --- | --- |
| **Gender** | 75 | |  | 0.102 |
| Male | 69 | 0.75 ± 0.11^(3)^ | |  |
| Female | 6 | | 0.10 ± 0.15 |  |
| **Age** | 75 | |  | 0.101 |
| ≥62^(4)^ | 39 | | 0.87 ± 0.16 |  |
| <62 | 36 | | 0.51 ± 0.12 |  |
| **TNM Stage** | 75 | |  | 0.808 |
| I | 28 | | 0.73 ± 0.17 |  |
| II/III | 35 | | 0.67 ± 0.15 |  |
| **Histological Grade** | 75 | |  | 0.447 |
| I/II  II-III/III | 55  20 | | 0.65 ± 0.12  0.84 ± 0.23 |  |

^1^RS: Reactive score

^2^Student's *t*-test; a *P* value less than 0.05 was considered statistically significant

^3^Data shown are mean ± s.e.m.

^4^The median age of the patients in this cohort was 62

**Supplementary Table 2. Summary of clinicopathological characteristics of the cohort of 92 lung adenocarcinoma patients**

| **Characteristics** | **Cases** | **PLANE abundance in Lung squamous cell carcinoma(RS^1^)** | | ***P* value^2^** |
| --- | --- | --- | --- | --- |
| **Gender** | 92 | |  | 0.325 |
| Male | 51 | | 0.34 ± 0.08^(3)^ |  |
| Female | 41 | | 0.48 ± 0.11 |  |
| **Age** | 92 | |  | 0.339 |
| ≥63^(4)^ | 48 | | 0.47 ± 0.11 |  |
| <63 | 44 | | 0.34 ± 0.04 |  |
| **TNM Stage** | 67 | |  | 0.276 |
| I /II | 41 | | 0.55 ± 0.12 |  |
| II/III | 26 | | 0.35 ± 0.10 |  |
| **Histological Grade** | 92 | |  | 0.143 |
| I/II  III | 57  35 | | 0.33 ± 0.07  0.54 ± 0.13 |  |

^1^RS: Reactive score

^2^Student's *t*-test; a *P* value less than 0.05 was considered statistically significant

^3^Data shown are mean ± s.e.m.

^4^The median age of the patients in this cohort was 63

**Supplementary Table 3. Summary of clinicopathological characteristics of the cohort of 89 colon cancer patients**

| **Characteristics** | **Cases** | **PLANE abundance in Lung squamous cell carcinoma(RS^1^)** | | ***P* value^2^** |
| --- | --- | --- | --- | --- |
| **Gender** | 89 | |  | 0.239 |
| Male | 51 | 0.43 ± 0.08^(3)^ | |  |
| Female | 38 | | 0.61 ± 0.12 |  |
| **Age** | 89 | |  | 0.203 |
| ≥68^(4)^ | 42 | | 0.41 ± 0.09 |  |
| <68 | 47 | | 0.66 ± 0.11 |  |
| **TNM Stage** | 80 | |  | 0.054 |
| I/II | 43 | | 0.66 ± 0.11 |  |
| III/IV | 37 | | 0.35 ± 0.10 |  |
| **Histological Grade** | 89 | |  | 0.750 |
| I/II | 72 | | 0.52 ± 0.07 |  |
| II-III/III | 17 | | 0.46 ± 0.21 |  |

^1^RS: Reactive score

^2^Student's *t*-test; a *P* value less than 0.05 was considered statistically significant

^3^Data shown are mean ± s.e.m.

^4^The median age of the patients in this cohort was 68

**Supplementary Table 4. Summary of proteins that interact with PLANE detected using mass spectrometry**

| No. | Entry name | Coverage [%] | MW [kDa] | Score |
| --- | --- | --- | --- | --- |
| 1 | HNRPM | 32 | 77.5 | 139.38 |
| 2 | M0QZM1 | 29 | 40 | 58.37 |
| 3 | K2C1 | 24 | 66 | 40.71 |
| 4 | K1C10 | 22 | 58.8 | 30.7 |
| 5 | K22E | 21 | 65.4 | 28.63 |
| 6 | NUCL | 11 | 76.6 | 19.43 |
| 7 | F5H5D3 | 17 | 57.7 | 18.9 |
| 8 | K1C9 | 15 | 62 | 18.85 |
| 9 | TBA1A | 20 | 50.1 | 18.12 |
| 10 | Q5ST81 | 16 | 41.7 | 12.98 |
| 11 | RFA1 | 10 | 68.1 | 12.31 |
| 12 | BIP | 7 | 72.3 | 9.96 |
| 13 | WDR76 | 6 | 69.7 | 8.81 |
| 14 | TBB2A | 14 | 49.9 | 8.8 |
| 15 | E7EWK3 | 4 | 91.4 | 8.17 |
| 16 | HNRPK | 9 | 50.9 | 7.68 |
| 17 | H0YIN9 | 16 | 22 | 7.62 |
| 18 | CSTF3 | 3 | 82.9 | 7.53 |
| 19 | E7EUT5 | 9 | 27.9 | 6.97 |
| 20 | DDX3X | 4 | 73.2 | 5.85 |
| 21 | PLOD1 | 2 | 83.5 | 5.85 |
| 22 | A6NLN1 | 7 | 56.5 | 5.39 |
| 23 | PCBP1 | 10 | 37.5 | 5.35 |
| 24 | IF4A1 | 8 | 46.1 | 4.87 |
| 25 | TOP2A | 2 | 174.3 | 4.71 |
| 26 | E9PKE3 | 5 | 68.8 | 4.65 |
| 27 | J3KTA4 | 4 | 69 | 4.61 |
| 28 | YBOX3 | 5 | 40.1 | 4.44 |
| 29 | D6RBZ0 | 7 | 35.7 | 4.37 |
| 30 | J3KT29 | 15 | 13.2 | 3.97 |
| 31 | HNRPF | 4 | 45.6 | 3.48 |
| 32 | M0R076N | 11 | 13.1 | 3.4 |
| 33 | API5 | 3 | 59 | 3.31 |
| 34 | DDX21 | 3 | 87.3 | 3.25 |
| 35 | WDR43 | 2 | 74.8 | 3.09 |
| 36 | H3BSS4 | 11 | 19.4 | 3.01 |
| 37 | K1C14 | 3 | 51.5 | 2.92 |
| 38 | M0R1V7 | 25 | 7.1 | 2.91 |
| 39 | I3L239 | 7 | 24.9 | 2.89 |
| 40 | F5GYT8 | 2 | 64 | 2.77 |
| 41 | RBM14 | 2 | 69.4 | 2.59 |
| 42 | E9PQD7 | 6 | 25.2 | 2.51 |
| 43 | A2A3R5 | 8 | 25 | 2.44 |
| 44 | H0Y449 | 5 | 42 | 2.4 |
| 45 | G3XAC6 | 3 | 48 | 2.39 |
| 46 | NOG1 | 2 | 73.9 | 2.34 |
| 47 | TGFB3 | 3 | 47.3 | 2.33 |
| **Supplementary Table 4, continued** | | | | |
| 48 | RS8 | 7 | 24.2 | 2.3 |
| 49 | PURB | 4 | 33.2 | 2.29 |
| 50 | D6RG13 | 8 | 25.6 | 2.25 |
| 51 | A0A087WVQ9 | 3 | 47.9 | 2.21 |
| 52 | RS23 | 8 | 15.8 | 2.19 |
| 53 | B5MCT8 | 6 | 16.6 | 2.17 |
| 54 | ALBU | 2 | 69.3 | 2.17 |
| 55 | FILA2 | 1 | 247.9 | 2.14 |
| 56 | B7ZM68 | 1 | 129.4 | 2.14 |
| 57 | F8WA26 | 19 | 18.5 | 0 |
| 58 | BROMI | 3 | 144.7 | 0 |
| 59 | AMD | 3 | 108.3 | 0 |

**Supplementary Table 5. Sequence similarity of transcripts in other species compared with human PLANE**

| Description | Total score | Query cover | E value |
| --- | --- | --- | --- |
| Homo sapiens | 1716 | 100% | 0.0 |
| Gorillas | 1467 | 92% | 0.0 |
| Mus musculus | 122 | 7% | 2e-12 |

**Supplementary Table 6. List of cell lines**

| **Cell lines** | **Source** | **Catalogue No.** |
| --- | --- | --- |
| A549 | ATCC | CCL-185 |
| MCF-7 | ATCC | HTB-22 |
| HCT116 | ATCC | CCL-247 |
| H1299 | National Science and Technology Infrastructure (NSTI, Shanghai, China) | SCSP-589 |
| CCC-HSF-1 | National Infrastructure of Cell Line Resource (NICR, Beijing, China) | 3111C0001CCC000069 |
| CCC-HIE-2 | National Infrastructure of Cell Line Resource (NICR, Beijing, China) | 3111C0001CCC000178 |
| Eca109 | Dr Xiao Ying Liu (Translational Research Institute, Henan Provincial People’s Hospital and People’s Hospital of Zhengzhou University, Zhengzhou, China) | N/A |
| NCI-H1975 | Prof Xiaoju Zhang (Respiration Department, Henan Provincial People’s Hospital, Zhengzhou, China) | N/A |
| NCI-H226 | Prof Xiaoju Zhang (Respiration Department, Henan Provincial People’s Hospital, Zhengzhou, China) | N/A |

**Supplementary Table 7. List of antibodies**

| Antibody (Ab) | Catalogue No. | Company | Dilution |
| --- | --- | --- | --- |
| E2F1 Rabbit mAb | ab179445 | Abcam (Cambridge, UK) | 1:1000 |
| H3K4me3 Rabbit mAb | ab1012 | Abcam (Cambridge, UK) | 1:500 |
| H3K27me3 Rabbit mAb | ab192985 | Abcam (Cambridge, UK) | 1:1000 |
| Normal mouse IgG | sc-2025 | Santa Cruz Biotechnology (Dallas, TX) | 1: 500 |
| Normal rabbit IgG mAb | ab172730 | Abcam (Cambridge, UK) | 1:1000 |
| HRP Conjugated AffiniPure Goat Anti-mouse IgG (H+L) | BA1050 | BOSTER Biological Technology (Wuhan, Hubei, P.R.C) | 1:2000 |
| HRP Conjugated AffiniPure Goat Anti-rabbit IgG (H+L) | BA1054 | BOSTER Biological Technology (Wuhan, Hubei, P.R.C) | 1:2000 |
| hnRNPM Rabbit pAb | 26897-1-AP | Proteintech Group (Wuhan, Hubei, P.R.C) | 1:500 |
| SC35 Mouse mAb | ab11826 | Abcam (Cambridge, UK) | 1:500 |
| CY3 Conjugated AffiniPure Goat Anti-rabbit IgG (H+L) | BA1032 | BOSTER Biological Technology (Wuhan, Hubei, P.R.C) | 1:200 |
| Chicken anti-Mouse IgG (H+L) Cross-Adsorbed Secondary Antibody, Alexa Fluor 488 | A21200 | Thermo Fisher Scientific (Waltham, MA) | 1:1000 |
| U1-70K Rabbit pAb | ab51266 | Abcam (Cambridge, UK) | 1:2000 |
| hnRNPK Rabbit mAb | ab52600 | Abcam (Cambridge, UK) | 1:10000 |
| NCOR2 Rabbit mAb | #62370 | Cell Signaling Technology (Danvers, MA)) | 1:1000 |

mAb: monoclonal antibody

pAb: polyclonal antibody

**Supplementary Table 8. List of reagents**

| **Reagent** | **Catalogue No.** | **Company** |
| --- | --- | --- |
| Doxycline | D9891 | Sigma-Aldrich (Saint Louis, MO) |
| Piece^TM^ RIPA Buffer | 89900 | Thermo Fisher Scientific (Waltham, MA) |
| Piece^®^ IP Lysis Buffer | 87787 | Thermo Fisher Scientific (Waltham, MA) |
| Cell counting kit 8 | HY-K0301 | Med Chem Express (Monmouth Junction, NJ) |
| Protease Inhibitor Cocktail | HY-K0010 | Med Chem Express (Monmouth Junction, NJ) |
| RiboLock RNase Inhibitor | EO0384 | Thermo Fisher Scientific (Waltham, MA) |
| 4% paraformaldehyde | AR1068 | BOSTER Biological Technology (Wuhan, Hubei, P.R.C) |
| Opti-MEM^TM^ Reduced Serum Medium | 31985070 | Thermo Fisher Scientific (Waltham, MA) |
| DNA extraction buffer | P1012 | Solarbio Life Sciences (Beijing, P.R.C) |
| Protease K | P1120 | Solarbio Life Sciences (Beijing, P.R.C) |
| Anti-GFP mAb-Magnetic Agarose | D153-10 | Medical& Biological Laboratories (Sakae, Nagoya, Aichi,Japan) |
| TRIzol^TM^ Reagent | 15596018 | Thermo Fisher Scientific (Waltham, MA) |

**Supplementary Table 9. List of primers used for recombinant plasmid**

| pcDNA3.1(+)-PLANE | Forward: CGGGATCCGCTGTCCCACGCGCCGGGT |
| --- | --- |
|  | Reverse: AACCTCGAGTGCATGTTCTCCAGATGTCCT |
| pcDNA3.1(+)-PLANE-R | Forward1: CGGGATCCGCTGTCCCACGCGCCGGGT  Reverse1: CTGAAATGACGGTCTCGTTCTATC  Forward2: GATAGAACGAGACCGTCATTTCAG  Reverse2: AACCTCGAGTGCATGTTCTCCAGATGTCCT |
| pcDNA3.1(+)-PLANE-△DFO | Forward1: GGATCCGCTGTCCCACGCGCCGGGT |
|  | Reverse1: GCGCAGGTAGGACTGCTATTCAGAC  Forward2: CTGAATAGCAGTCCTACCTGCGCCC  Reverse2: CTCGAGTGCATGTTCTCCAGATGTCCT |
| pcDNA3.1(+)-PLANE-R-△331-751 | Forward1: GGGATCCGCTGTCCCACGCGCCGGGT  Reverse1: TGATATTCTGTGACTGGTTCAGACCCCTTCA  Forward2: GAAGGGGTCTGAACCAGTCACAGAATATCAG |
|  | Reverse2: CCTCGAGTGCATGTTCTCCAGATGTCCT |
| pEGFP-C1-hnRNPM | Forward: TAGGTACCATGGCGGCAGGGGTCGAAG  Reverse: CGGGATCCTTAAGCGTTTCTATCAATTCGAAC |
| pEGFP-C1-hnRNPM-△RRM1 | Forward1: TAGGTACCATGGCGGCAGGGGTCGAAG  Reverse1: TGTTCACCATCAGGCTGTATCTTTTAGT  Forward2: ACTAAAAGATACAGCCTGATGGTGAACA  Reverse2: CGGGATCCTTAAGCGTTTCTATCAATTCGAAC |
| pEGFP-C1-hnRNPM-△RRM2 | Forward1: TAGGTACCATGGCGGCAGGGGTCGAAG  Reverse1: TGGTAAGGCCCTCTCACTGTGCTTCCAAGTC  Forward2: GACTTGGAAGCACAGTGAGAGGGCCTTACCA  Reverse2: CGGGATCCTTAAGCGTTTCTATCAATTCGAAC |
| pEGFP-C1-hnRNPM-△RRM3 | Forward: TAGGTACCATGGCGGCAGGGGTCGAAG |
|  | Reverse: CGGGATCCTGGCAGGCCTTCCTGGCCA |

**Supplementary Table 10. List of qRT-PCR primers**

| PLANE | Forward: TACATACAGTGACCCAAAGAGCA |
| --- | --- |
|  | Reverse: CAGTGCTTCTGAACGCCTCTT |
| NCOR2 mRNA | Forward: GTACAAAGACCGCCAGGTCA |
|  | Reverse: TGATGCGATCAGGCCAAAGT |
| NCOR2 pre-mRNA | Forward: GGCTTCTTGGCCCATCT |
|  | Reverse: AAGGCTCCCTGACTCCC |
| GAPDH | Forward: GCTCTCTGCTCCTCCTGTTC  Reverse: ACGACCAAATCCGTTGACTC |
| β-actin | Forward: GGACTTCGAGCAAGAGATGG  Reverse: AGCACTGTGTTGGCGTACAG |
| U6 | Forward: TCGCTTCGGCAGCACATAT  Reverse: ATTTGCGTGTCATCCTTGC |
| MELTF | Forward: GAGCCCCCTGGAGAGATACT  Reverse: CATCGTCCTACGTGCTTCCT |

**Supplementary Table 11. List of RT-PCR primers and RNA pulldown probes**

| ChIP-E2F1-BR | Forward: AGGGGGGCAGGGCTAGTAG  Reverse: ACCTAGATCCTGCCTCCC |
| --- | --- |
| NCOR2-202/NCOR-001/-005 | Forward: GAACATGCCAGCACCAACA  Reverse: GGAATGGCGTGGAACCTG |
| ChIP-H3K4me3-BS/-H3K27me3-BS | Forward: AACGCGATTCCAGTGAGGT  Reverse: GCGACGCCGAGTTTCTTT |
| MELTF-AS1-201 | Forward: CTGCTGAGACGACATCCCTT  Reverse: ACGCCCAGCTCACCTGAT |
| MELTF-AS1-202 (PLANE) | Forward: TACATACAGTGACCCAAAGAGCA  Reverse: CAGTGCTTCTGAACGCCTCTT |
| MELTF-AS1-203 | Forward: GCAACCCACGCTTCGAG  Reverse: TCCTGCCTCCCAAGGTG |
| MELTF-AS1-204 | Forward: CTCCCCACAAACCTAAACA  Reverse: GTAGCCACAGAACGGTCAT |
| MELTF-AS1-205 | Forward: GCGGCGCCTCAGATGC  Reverse: GAGTGTGAACGCTCAAAACGG |
| dChIRP-PLANE | Forward: CTGGACTGCTGCGAACG  Reverse: TGGCTGGGGCTGGACTA |
| dChIRP-intron 45 | Forward: CTTGTGACTTTATTTTTGTGCGTGT  Reverse: CAAGGAGACAGATGGGCCAAG |
| RRBP1 | Forward: AGGTCCTGGAGGTGCCATTTC  Reverse: GCCAGGTGTCCTGAATGATGC |
| DNHD1 | Forward: GCCCATCTTTGATACCTTC  Reverse: CTAACCAGACCTCGTGCC |
| ADH6 | Forward: TGTAAAGCAGCAGGAGCA  Reverse: CAGCAACCAGTTTAGGGA |
| SLC25A4 | Forward: GCTCCTGCGTTGCTAAGACA  Reverse: ACAGGCATGAGCCACAGCA |
| PTPN4 | Forward: AGTAAGCCCTTGGCACGGA  Reverse: ACATTGAATCCAAACCTCCC |
| hnRNPM-BS | Forward: GGCTTCTTGGCCCATCT  Reverse: AAGGCTCCCTGACTCCC |
| lncCyt b | Forward: TTGTTTGATCCCGTTTCGTG  Reverse: ACATCGGCATTATCCTCCTG |
| PLANE-biotin-probes | AS1: ATGACTTGCTCTTTGGGTC  AS2: CATTCTAGTGATTCTGAGC  S1: GACCCAAAGAGCAAGTCAT  S2: GCTCAGAATCACTAGAATG |
| NCOR2 pre-mRNA-biotin-probes | AS1: TTGGACAACTGCAACTCTC  AS2: GTCTGCTGTTTGCAATAGC  AS3: TAAACTGTCTGCTGTTTGC  S1: GAGAGTTGCAGTTGTCCAA  S2: GCTATTGCAAACAGCAGAC  S3: GCAAACAGCAGACAGTTTA |
| PLANE (*in vitro* transcription) | Forward: TAATACGACTCACTATAGGGGCTGTCCCACGCGCCGGGT  Reverse:  TGCATGTTCTCCAGATGTCCTTTAT |
| **Supplementary Table 11, continued** |  |
| AS PLANE (*in vitro* transcription) | Forward: TAATACGACTCACTATAGGGTGCATGTTCTCCAGA  TGTCCTTT  Reverse: GCTGTCCCACGCGCCGGGTCCC |
| PLANE △DFO (*in vitro* transcription) | Forward1: TAATACGACTCACTATAGGGGCTGTCCCACGCGCCGGGT  Reverse1:  GCGCAGGTAGGACTGCTATTCAGAC  Forward2: CTGAATAGCAGTCCTACCTGCGCCC  Reverse2: TGCATGTTCTCCAGATGTCCTTTAT |
| PLANE 1-330 (i*n vitro* transcription) | Forward: TAATACGACTCACTATAGGGGCTGTCCCACGCGCCGGGT  Reverse: TTCAGACCCCTTCACCCCAGAAC |
| PLANE 331-751 (*in vitro* transcription) | Forward: TAATACGACTCACTATAGGGGTTCTGGGGTGAAGGGGTCTG  Reverse: GTTGCAGAACACAAGTCCCTCTCGA |
| PLANE 752-951 (*in vitro* transcription) | Forward: TAATACGACTCACTATAGGGCAGTCACAGAATATCAGGTGAGC  Reverse: TGCATGTTCTCCAGATGTCCTTTAT |
| NCOR2 intron45 (*in vitro* transcription) | Forward: TAATACGACTCACTATAGGGTCTGTCTGTCTGTCTCTCTCTC  Reverse: CTGCAGGGGGACAAGATGGG |
| NCOR2 intron47 (*in vitro* transcription) | Forward: TAATACGACTCACTATAGGGTCAGGTCCCAGCGAGCCA  Reverse: GGAGTATAATTCGCTTTTTAATTAG |
| NCOR2-intron 45-△PLANE-BR (*In vitro* transcription) | Forward1: TAATACGACTCACTATAGGGTCTGTCTGTCTGTCTCTCTCTC  Reverse1: GGCGGACAGCAGTGTGAGTGTGGGCAGGAGGGC  Forward2:  GCCCTCCTGCCC ACACTCACACTGCTGTCCGCC  Reverse2: CTGCAGGGGGACAAGATGGG |

**Supplementary Table 12. List of siRNAs/shRNAs**

| PLANE | siRNA1/shRNA.1: GACCCAAAGAGCAAGUCAU |
| --- | --- |
|  | siRNA2/shRNA.2: GCUCAGAAUCACUAGAAUG |
| NCOR2 | siRNA: UGGUGGAGGAUGAGGAGAU |
| hnRNPM | siRNA.1: GCAUCGGAAUGGGAAACAU |
|  | siRNA.2: CCAUUUGACUGUUUGCAUU |
| MELTF | siRNA.1: GGAUGGAGGAGCCAUCUAU  siRNA.2: GGGCGAAGUGUACGAUCAA |
